# Leptomeningeal collaterals regulate reperfusion in ischemic stroke

**DOI:** 10.1101/2023.02.25.529915

**Authors:** Nadine Felizitas Binder, Mohamad El Amki, Chaim Glück, William Middleham, Anna Maria Reuss, Adrien Bertolo, Patrick Thurner, Thomas Deffieux, Hannah-Lea Handelsmann, Philipp Baumgartner, Cyrille Orset, Philipp Bethge, Zsolt Kulcsar, Adriano Aguzzi, Mickael Tanter, Denis Vivien, Matthias T. Wyss, Andreas Luft, Michael Weller, Bruno Weber, Susanne Wegener

## Abstract

Recanalization is the mainstay of ischemic stroke treatment. However, even with timely clot removal, many stroke patients recover poorly. Leptomeningeal collaterals (LMCs) are pial anastomotic vessels with yet unknown functions. Utilizing a thrombin-based mouse model of stroke and the gold standard fibrinolytic treatment rt-PA, we here show that LMCs play a critical role in preserving vascular function in ischemic territories. We applied laser speckle contrast imaging, ultrafast ultrasound, and two-photon microscopy, to show that after thrombolysis, LMCs allow for gradual reperfusion resulting in small infarcts. On the contrary, in mice with poor LMCs, distal segments of recanalized arteries collapse and deleterious hyperemia causes hemorrhage and mortality. Accordingly, in stroke patients with poor collaterals undergoing thrombectomy, rapid reperfusion resulted in hemorrhagic transformation and unfavorable recovery. Thus, we identify LMCs as key components regulating reperfusion after stroke. Future therapeutic interventions should aim to enhance collateral function, allowing for gradual reperfusion of ischemic tissues after stroke.

## Main

Ischemic stroke, caused by the abrupt blockage of a brain-feeding artery, leads to disability and death in millions of people every year ^1^. Current ischemic stroke treatments aim at restoring blood flow by either intravenous thrombolysis, mechanical thrombectomy, or a combination of both ^2^. However, despite timely and successful recanalization of the occluded vessel, many patients show insufficient clinical improvement. This has been referred to as “futile recanalization” ^3, 4^. Recanalization is the prerequisite to establish reperfusion, i.e. recovery of blood flow in the brain. Yet, several processes including distal clot fragmentation ^5^, pericyte constriction ^6^, or neutrophil obstruction of capillaries ^7^ may hamper reperfusion of the ischemic brain. Therefore, progressive infarct expansion in futile recanalization is caused by reperfusion failure ^8^ .

Leptomeningeal collaterals (LMCs) are pial anastomotic vessels connecting a fraction of the terminal branches of the middle cerebral artery (MCA) with terminal branches of the anterior (ACA) and posterior (PCA) cerebral arteries ^9^. Although LMCs exhibit minimal flow under normal physiological conditions, when obstruction occurs in a major supply artery, flow across them is recruited and provides partial support of cerebral blood flow (CBF) in the ischemic core and peri-infarct region ^9, 10^. Recruitment of collateral flow is achieved by vasodilation and existence of a pressure gradient that results in redistribution of flow from neighboring areas to the ischemic tissue ^11^. The extent of LMCs varies in humans and rodents ^12^. In stroke patients, extensive LMCs are associated with better outcome after thrombolysis and mechanical thrombectomy ^13–15^. While LMCs alleviate the severity of ischemic injury, little is known about how differences in collateral extent affect reperfusion after recanalization. In the present study, we hypothesized that the presence of abundant LMCs provides hemodynamic support prior to and during reperfusion, thus preventing futile recanalization after stroke therapies.

We used a thrombin model of stroke and thrombolysis in three mouse strains with significant differences in collateral extent. We monitored reperfusion processes *in vivo* to reveal the contributions of LMCs to vessel integrity as well as spatial and temporal dynamics of tissue reperfusion. We then extended these findings in mice to patients with acute ischemic stroke that received thrombectomy, and provide direct translational evidence in support of a protective role of LMCs during reperfusion.

## Results

### Variation in leptomeningeal collaterals in selected mouse strains

To contrast arterial networks with different extents of LMCs, we examined C57BL/6, Rabep2-/-, and Balb-C mice ^16^. Rabep2-/- mice have a C57BL/6 background but, due to a disruption of the Rabep2 gene previously shown to be important in LMC formation during development, have a reduced number of LMCs ^16, 17^. For 3D vessel visualization and LMC quantification, we performed brain clearing and immunostaining based on iDISCO ^18^. We imaged smooth muscle actin (α-SMA)-positive cells throughout the entire brain using light sheet microscopy (Fig. 1a,b, Supplementary Fig. 1) ^19^. LMCs were defined as α-SMA-positive vessels anastomosing opposing distal branches of the MCA and ACA. Consistent with previous reports ^16, 21^, we observed an average of approximately 10 MCA-ACA collaterals per hemisphere in C57BL/6 and 2-3 in Rabep2-/- mice, and none in most (occasionally 1) Balb-C mice (Fig. 1c). Thus, we denoted LMCs as “abundant” in C57BL/6, “intermediate-to-poor” in abundance in Rabep2-/-, and “poor” in Balb-C animals.

**Fig. 1:**
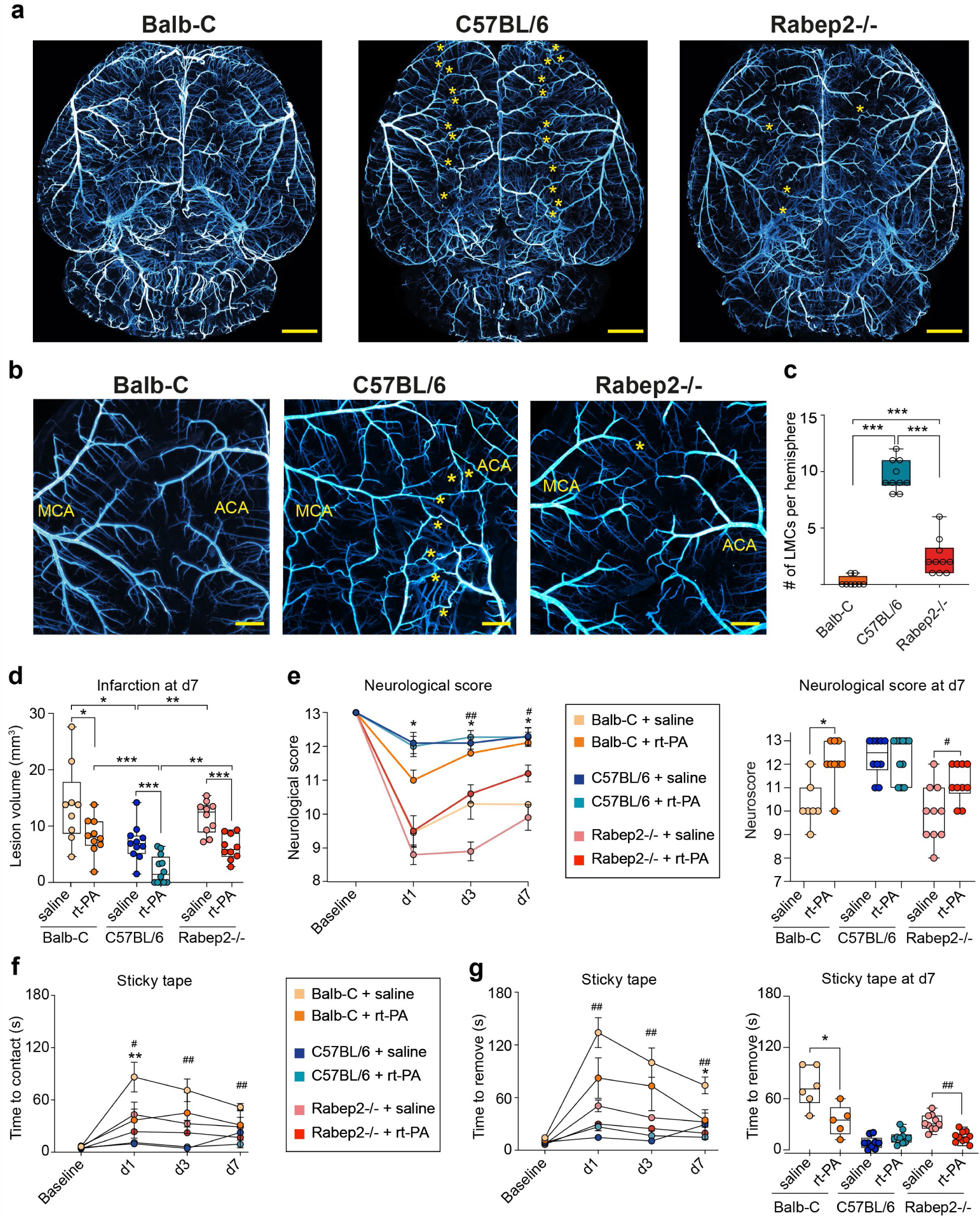
Quantification of LMCs in Balb-C, C57BL/6 and Rabep2-/-. Infarct volume and sensorimotor function in mice with different LMCs (**a**) Representative overview images of iDISCO cleared and anti-SMA-Cy3 stained brains of Balb-C, C57BL/6 and Rabep2-/- mice. Yellow stars indicate LMCs. Scale bar, 1cm. (**b**) Close- up images of watershed area, yellow stars indicate LMCs. Scale bar, 200µm. (**c**) Quantification of LMCs per hemisphere in all three strains (n=5 per strain). (**d**) Bar graph depicting infarct volumes from individual mice from the three different strains at day 7, ∗p < 0.05, ∗∗∗p < 0.001 in two-tailed Mann-Whitney U-test. (**e**)- (**g**) Neurological score and sticky tape removal assessment in saline and rt-PA-treated mice at days 1, 3, and 7 after stroke. (n = 10); ∗ Balb-C saline vs rt-PA, # Rabep2-/- saline vs rt-PA; ∗/ # p < 0.05, ∗∗/##p < 0.01; in two-tailed Mann- Whitney U-test.

### Thrombolysis-induced MCA recanalization is independent of LMC extent

Next, we sought to determine whether the presence of LMCs impact clot formation and the dissolution processes during thrombolysis. We employed the thrombin model of stroke in C57BL/6, Rabep2-/-, and Balb-C mice ^20, 21^ (Supplementary Figs. 2, 3). This model closely mimics the clinical situation of stroke patients with natural clot formation and rt-PA-based intravenous thrombolysis, allowing for direct *in vivo* visualization of reperfusion. We investigated the formation of the clot after thrombin microinjection and clot presence at 2 hours post stroke (90 minutes after the start of rt-PA or vehicle infusion). Independent of strain, thrombin injection led to an occlusion of the MCA at its M2 bifurcation, which persisted for 2 hours in 60-75% of the control treated mice. Rt-PA infusion at 30 minutes after stroke induction led to complete or partial MCA-M2 recanalization (Balb-C: 94.1%; C57BL/6: 95.3%; Rabep2-/-: 92.9%) independent of LMC status (Supplementary Fig. 2). Thus, we did not observe that more abundant LMCs augmented clot dissolution on thrombolytic treatments and enhanced recanalization.

### Stroke outcome depends on number of LMCs

We then tested whether abundant LMCs preserve tissue integrity and neurological function in stroke. Mice receiving either rt-PA or vehicle were monitored for seven days after induction of stroke. Infarct volumes on day 7 were largest in Balb-C, intermediate in Rabep2-/-, and smallest in C57BL/6 mice (Fig. 1 d-g), confirming previous studies employing permanent distal M1-MCA ligation wherein robust collateral circulation limited infarct size ^17^. Thrombolysis significantly reduced ischemic tissue damage in all three strains compared to control treatment (Balb-C: 13.84 ± 2.34 mm^3^ vs. 8.22 ± 1.01 mm^3^; Rabep2-/-: 11.59 ± 0.91 mm^3^ vs. 6.24 ± 0.70mm^3^; C57BL/6: 7.07 ± 0.96 mm^3^ vs. 2.26 ± 0.72 mm^3^) (Fig. 1d). This was reflected in better sensorimotor function after rt-PA in Balb-C and Rabep2-/-, but not in C57BL/6 mice, which showed only minimal deficits overall (Fig. 1e). In general, sensorimotor functions assessed by the sticky tape test and a composite neurological score were better for C57BL/6 mice, which having abundant LMCs experienced only minimal deficits (Fig. 1e-g).

### LMCs are recruited during stroke and reperfusion

To examine the contribution of LMCs to stroke and reperfusion, we performed *in vivo* two-photon imaging in the MCA-ACA watershed area of C57BL/6 mice (Fig. 2). Immediately after stroke, LMCs underwent dilation and collateral flow was recruited from the ACA to the MCA territory (Fig. 2d, 2e). Following recanalization of the MCA, collateral diameter and flow returned to baseline values, demonstrating the contribution of LMCs to blood flow towards the occluded territory. Therefore, LMCs are indeed recruited during stroke, augmenting flow towards the affected territory.

**Fig. 2:**
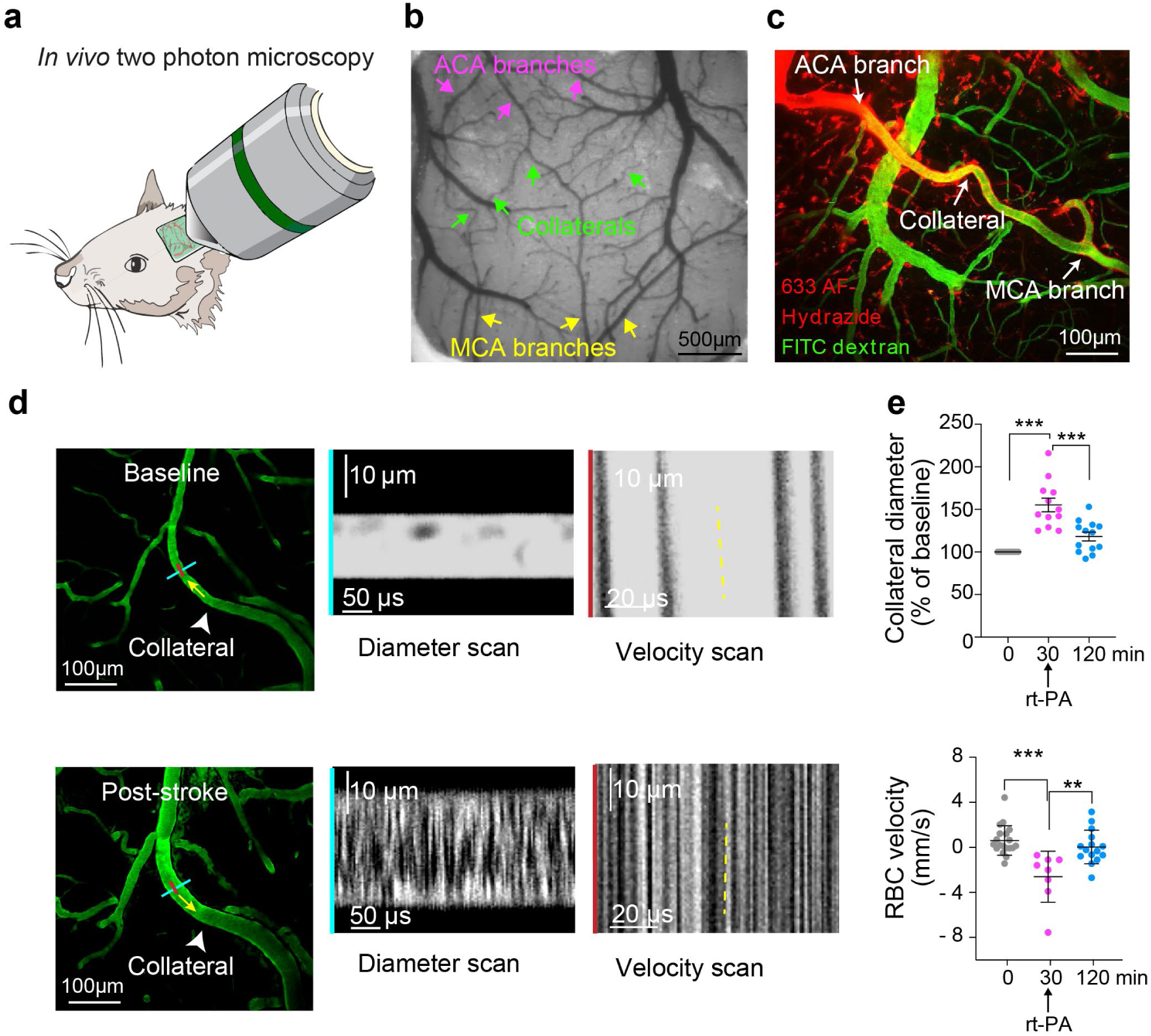
Two-photon imaging of LMCs pre- and post-stroke in C57BL/6 mice. (**a**) Schematic drawing of two-photon setup. (**b**) Cranial window of C57BL/6 mouse. This strain was used here because it has extensive LMCs. Arrowheads displaying MCA branches and LMCs. (**c**) Image during two-photon acquisition. Blood plasma is labelled in green using FITC dextran. To differentiate arteries from veins, 633 Alexa Fluor (AF)- Hydrazide (red) staining was used (arterial vessel wall in red). (**d**) On the left, representative image of LMC response to stroke in C57BL/6 mice. In the middle, kymographs of LMC diameter changes post-stroke, generated by transverse line scans (blue lines in left images). On the right, kymographs of red blood cell flow in collaterals in baseline and post-stroke generated by line scans parallel to the flow (red lines in left images). Dark streaks represent red blood cells, grey streaks represent the fluorescent tracer-filled LMC lumen. (**e**) LMC RBC velocity and diameter changes in C57BL/6 at baseline (0), 30 and 120min after stroke induction; ∗∗p < 0.01; ∗∗∗p < 0.001 two-tailed t test, n=5.

### LMCs affect the dynamics of blood flow recovery during reperfusion

Having documented recruitment of LMCs during ischemia and reperfusion at the single-vessel- evel, we next explored the effects of LMC recruitment on recovery of blood flow to the ischemic territory during thrombolysis. Laser speckle contrast imaging (LSCI) was used to provide continuous wide-field perfusion assessment with high temporal resolution and spatial resolution at a depth of 200 μm ^22^ at baseline, during, and for 120 min after thrombin occlusion (Fig. 3a, 3b). After occlusion of the MCA, all three strains evidenced a significant drop in perfusion of the distal MCA territory (M4/M5 territory) to 20-30% of baseline (Fig. 3b, c). Cortical perfusion remained low for 120 min after induction of stroke followed by sham-thrombolysis (Supplementary Figs. 4, 5). Significant reperfusion followed rt-PA administration in Balb-C and Rabep2-/- mice (60.2 ± 3.77% and 59.2 ± 3.66% of baseline). Surprisingly, in C57BL/6 mice with abundant collaterals and irrespective of rt-PA administration, CBF recovered more slowly and to only about 50% of baseline (47.3 ± 4.27% vs. 47.7 ± 4.53%) (Fig. 3b).

**Fig. 3:**
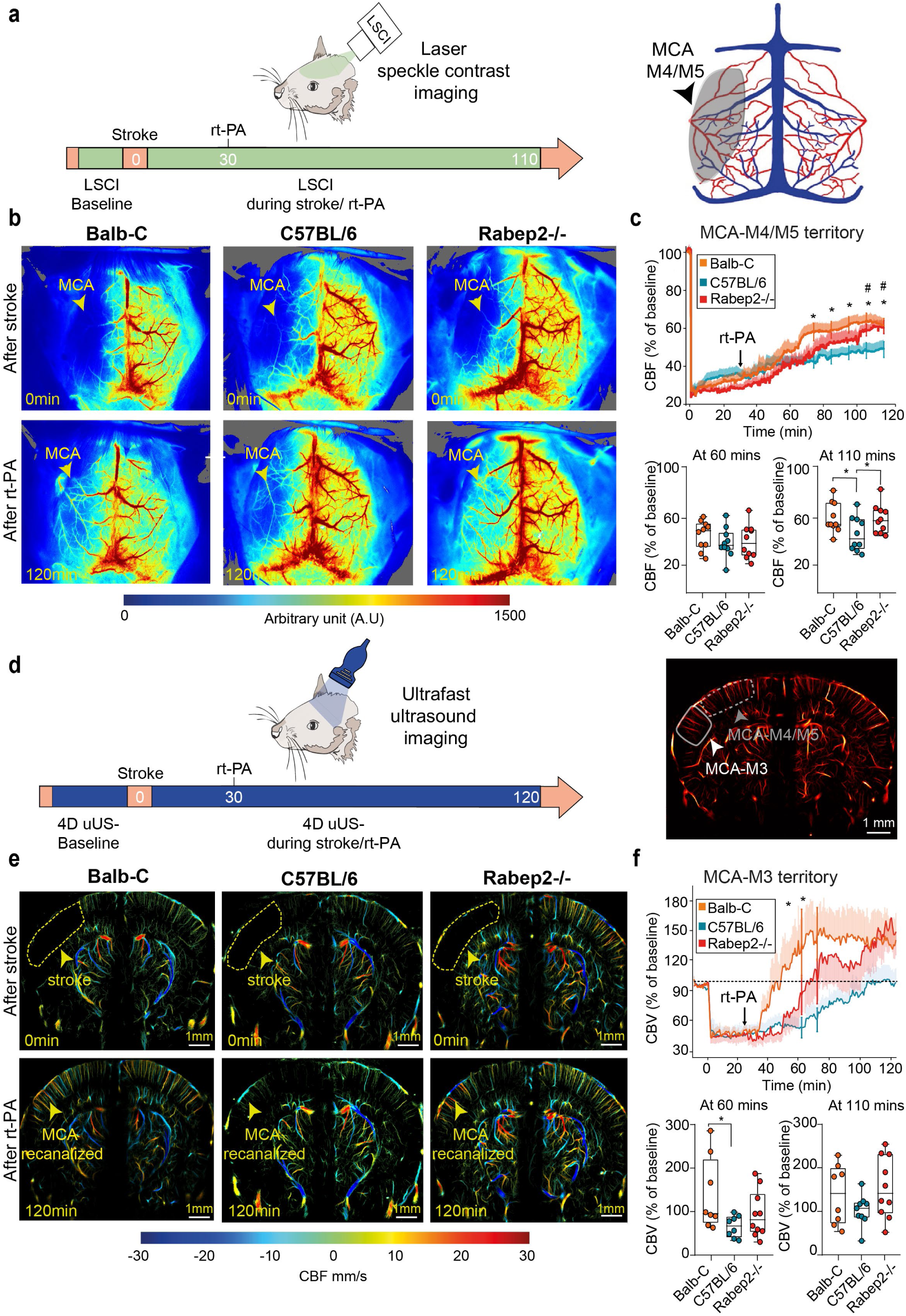
Reperfusion after thrombin occlusion and thrombolysis assessed by LSCI and uUS. (**a**) Experimental setup, containing 110 minutes post-stroke CBF monitoring (top). Schematic drawing of the LSCI view on pial vessels and the MCA-M4/M5 region of interest (ROI) used for analyses (bottom). (**b**) Representative LSCI images showing cortical perfusion after stroke induction (upper row) and thrombolysis (bottom row) of Balb-C, C57BL/6 and Rabep2-/- mice at 2 hours after stroke. Arrowhead indicating perfusion status of MCA branch. (**c**) LSCI recordings for the MCA-M4/M5 ROI compared to baseline in rt-PA treated Balb-C, C57BL/6, and Rabep2-/- mice, n = 10/group, ∗p < 0.05, two-tailed Mann-Whitney U-test. (**d**) Experimental setup of ultrafast ultrasound (uUS), containing 2 hours post-stroke CBV monitoring (top). 3D ROI selection of the MCA-M3 territory in view of coronal section (bottom). (**e**) Representative 2D ULM acquisitions immediately after stroke and 90 minutes after rt-PA start in Balb-C, Rabep2-/- and C57BL/6 mice. (**f**) 4D ultrasensitive Doppler analysis of rt-PA treated Balb-C (n=8), Rabep2-/- (n=10) and C57BL/6 (n=9) animals of the MCA-M3 territory. ∗p < 0.05, two-tailed Mann-Whitney U-test

Based on the above discrepancy between slower and less cortical reperfusion in the MCA- M4/M5 territory, yet excellent functional and tissue outcome in C57BL/6 mice (abundant collaterals), we hypothesized that LMCs facilitate reperfusion of more proximal territory in C57BL/6 mice that is not accessible to LSCI. Therefore, we next used ultrafast ultrasound (uUS) with 4D ultrasensitive Doppler recordings and ultrasound localization microscopy (ULM) ^23^ to investigate the more proximal MCA-supplied areas, just distal to the site of thrombin microinjection and clot formation.

We next applied uUS to measure cerebral blood volume (CBV) in deep (8 mm maximum) tissue during stroke and thrombolysis for 2 hours (Fig. 3d-f). We first evaluated CBV in the MCA- M4/M5 territory that had been assessed with LSCI in the previous experiment in Fig. 2. Consistent with the LSCI findings, thrombolysis resulted in higher reperfusion levels within the area supplied by the MCA-M4/M5 in mice with poor and poor-to-intermediate LMCs (Balb-C, Rabep2-/-), compared to animals with abundant LMCs (C57BL/6) (Supplementary Fig. 6). In agreement with our previous LSCI results, complete reperfusion was not achieved in this superficial cortical region. However, in more proximal MCA-M3 territory, thrombolysis resulted in reperfusion to, or exceeding 100% of baseline in all three strains (Fig. 3f), indicating that recanalization-induced reperfusion of proximal and distal MCA-supplied brain areas varied greatly depending on LMC status: mice with poor or poor-to-intermediate LMCs showed hyperaemic (above 100% baseline) reperfusion in the affected M3 territory (Balb-C 138.8 ± 23.1%, Rabep2-/- 145.8 ± 20.3%) (Fig. 3f). Furthermore, the time to reach pre-stroke baseline perfusion levels after thrombolysis was also notably different: mice with abundant LMCs underwent gradual reperfusion, reaching 100% of pre-stroke values in 98.5 ±7.77 minutes. In contrast, mice with poor or poor-to-intermediate LMCs underwent significantly earlier (Balb- C) and faster/steeper rates of reperfusion (Balb-C and Rabep2-/-), reaching 100% at 66.37 ± 12.59 (Balb-C) or 83.7 ± 9.54 minutes (Rabep2-/-), respectively.

### Rapid reperfusion and hyperemia in mice with poor collaterals is associated with loss of vascular tone in distal MCA branch segments

To reveal the structural and hemodynamic underpinnings of the observed different reperfusion responses, we used intravital two-photon microscopy (Fig. 4). We analyzed diameter and blood flow in the distal MCA branches (MCA-M5 segment) in Balb-C, C57BL/6, and Rabep2-/- mice during stroke and reperfusion. Following MCA occlusion, all animals sustained a large decrease in RBC flow velocity in the MCA-M5 segment (Fig. 4c). The drop in blood flow was less severe in C57BL/6 mice (abundant collaterals), suggesting that abundant LMCs maintain residual perfusion to the outer MCA territory after occlusion. Interestingly, the diameter of M5 segment arterioles as assessed between 20 and 30 min after thrombin occlusion declined significantly in Balb-C mice with poor collaterals, while reduction in diameter was much less in Rabep2-/- and absent in C57BL/6 mice (Fig. 4b,c).

**Fig. 4:**
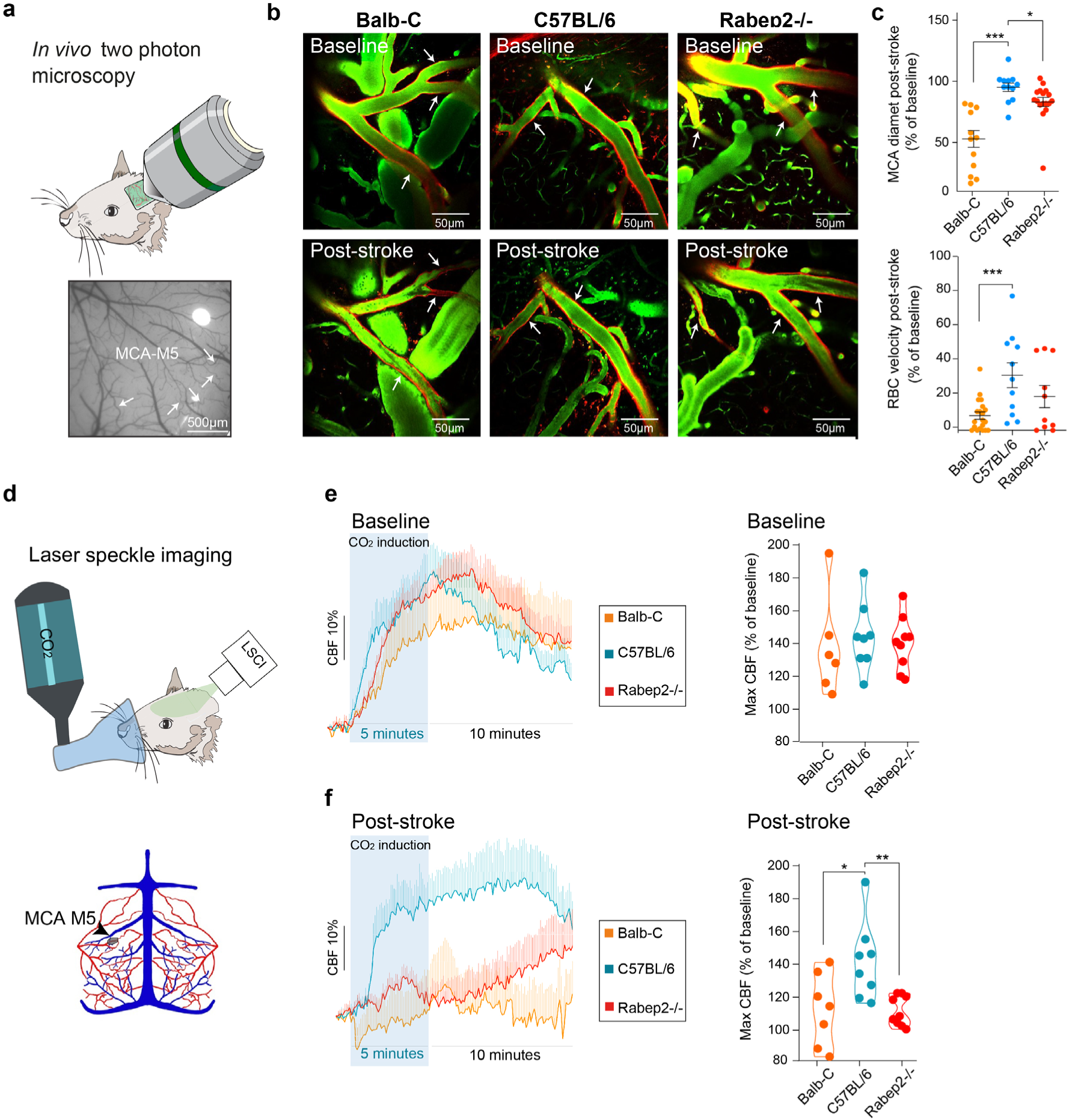
Hypercapnic challenge to probe vascular function before and after stroke. (a) Cranial window of a C57BL/6 mouse. White arrowheads displaying MCA M5 segments. (b) Representative images of distal MCA branches (white arrowheads) of Balb-C, C57BL/6 and Rabep2-/- mice pre- and post- stroke. (**c**) Quantification of RBC velocity and distal MCA diameter post-stroke (taken between 20-30 minutes) compared to the pre-stroke measurement (% change) is shown in n = 5 mice per strain; ∗p < 0.05; two-tailed t test. (**d**). Schematic drawing of hypercapnia setup (upper panel) and of the LSCI view on pial vessels and the MCA-M5 region of interest (ROI) used for analyses (lower panel). (**e-f**) CBF changes before, during, and after hypercapnia during baseline (**e**) and 30 minutes after thrombolysis (**f**). Violin plots indicate maximal CBF increase during 15 minutes after CO_2_ start. In n=7 Balb-C, n=8 C57BL/6, n=8 Rabep2-/-; ∗∗p < 0.001; ∗∗∗p < 0.001, two-tailed Mann-Whitney U-test

To test whether the decline in diameter reflected loss of vessel function, we assessed cerebrovascular reactivity. During pre-stroke and post-thrombolysis conditions, mice received a 5 min CO_2_ challenge via facemask, while we measured perfusion using LSCI (Fig. 4d). During baseline, hypercapnia induced an increase in CBF to 137.3 ± 10.64% (Balb-C), 144.1 ± 7.31% (C57BL/6) and 140 ± 5.45% (Rabep2-/-) within the MCA-M5 supplied territory (Fig. 4e). However, when challenged 60 minutes after thrombin microinjection and 30 minutes after start of rt-PA infusion, the hyperemic response was attenuated in Balb-C to 113.4 ± 8.94% and Rabep2-/- to 111.3 ± 3.02%, while preserved in C57BL/6 mice (141.5 ± 8.44%) (Fig. 4f). These data indicate a loss of vascular regulation capacity in distal blood vessels after clot removal in mice with poor and poor-intermediate LMCs.

### Rapid hyperemia in mice with poor collaterals associates with intracerebral hemorrhage and poor functional outcome

Given that intracerebral hemorrhage (ICH) is a major complication of recanalization treatments after stroke, we asked whether the hyperemic reperfusion in mice with poor LMCs was associated with tissue injury. Therefore, we analyzed mortality and occurrence of ICH for up to seven days after stroke. 40% of all rt-PA treated Balb-C mice died in the subacute phase after stroke (days 1 to 4), while no mortality occurred in Rabep2-/- or C57BL/6 mice (Fig. 5a, 5b). LSCI analysis of the MCA-M4/M5 territory revealed that animals that died prematurely had significantly faster reperfusion that reached higher values in the initial two hours after stroke, compared to the ones surviving (Fig. 5c). Notably, even within the group of Balb-C animals (poor LMCs) with rt-PA thrombolysis, significantly higher reperfusion levels were reached in animals that died versus those that survived (Supplementary Fig. 7a). Furthermore, all rt-PA treated Balb-C and a subset of Rabep2-/- treated animals showed evidence of ICHs after reperfusion, while none of the C57BL/6 mice did (Fig. 5d, Supplementary Fig. 7b). These data demonstrate that a fast and steep reperfusion in cortical areas supplied by the MCA-M4/M5 segment was associated with a higher risk of complications (ICH/death).

**Fig. 5:**
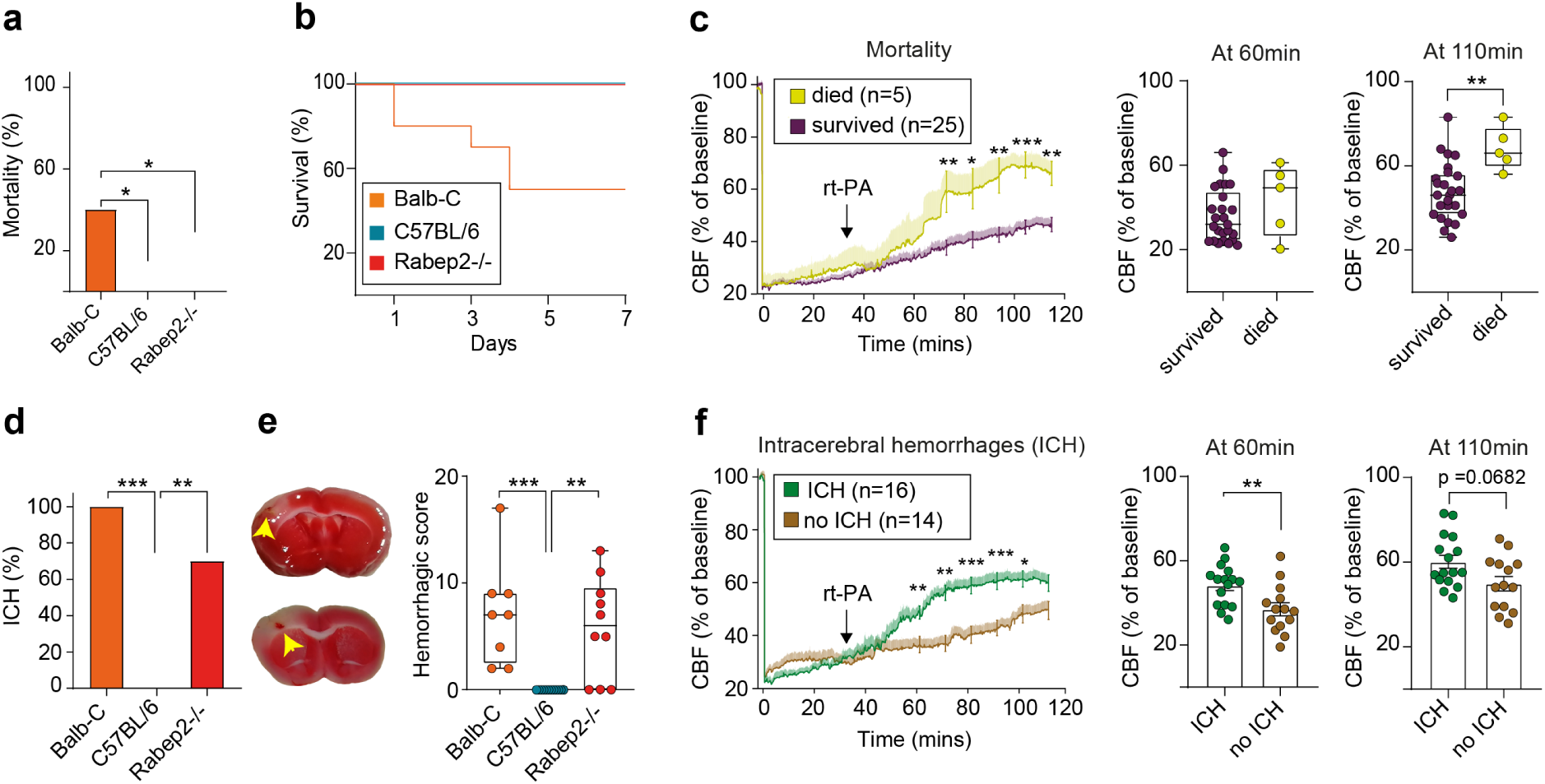
Mortality and intracranial hemorrhage in Balb-C, C57BL/6 and Rabep2-/- after thrombolysis. (**a**) Percentage of mortality within 7 days in rt-PA treated Balb-C, C57BL/6 and Rabep2-/- mice (n=10/strain) and (**b**) survival curve. (**c**) Sub-grouped LSCI analysis of animals, which died vs. survived of all strains receiving rt-PA. (**d**) – (**f**) Intracranial hemorrhage (ICH) assessment. (**d**) Overall percentage of rt-PA treated animals showing ICHs plus representative TTC image on d7. (**e**) Quantifications of hemorrhages (yellow arrows). The hemorrhagic score contains microscopic and macroscopic assessment.^7^ (**f**) Sub-grouping of LSCIs of rt-PA treated mice that showed ICHs vs mice without.

### Early venous filling in stroke patients undergoing thrombectomy indicates unfavorable outcome

Finally, we explored whether the observed reperfusion patterns in mice with different extents of LMCs and their association with stroke treatment outcome could be of relevance for stroke patients. The only direct assessment of reperfusion after stroke treatment is the immediate post-interventional DSA from patients with large vessel occlusion (LVO) stroke undergoing thrombectomy. We examined whether rapid reperfusion after recanalization of a large vessel occurred in stroke patients associated with poor collateral status and adverse outcomes. Out of 96 patients undergoing intra-arterial treatment for ischemic stroke at the University Hospital Zurich Stroke Center between 01-07/2021, we included data of 33 patients with distal MCA- M1 or M2 occlusion who had successful recanalization (modified treatment in cerebral infarction, mTICI ^24^ grade 2b or 3. We excluded patients with additional ipsilateral carotid artery stenosis or occlusion, distal clot embolism, without clinical follow-up according to the modified Rankin Scale (mRS) at 3 months, and patients who declined use of their data for research (see patient flow chart on Supplementary Fig. 8). Post-thrombectomy data from DSA were color-coded according to the time of contrast filling (in seconds; Fig. 6). From these time-coded DSA series, two senior neuroradiologists blinded to patient outcome or follow-up imaging categorized patients into those with early venous filling (n =23) and without early venous filling (n=10) after thrombectomy, approximating the rapid versus slow reperfusion pattern observed in the rodent stroke model. The two groups were then compared regarding clinical characteristics (age, sex), collateral score (poor, moderate, good based on pre-thrombectomy CT angiography data), initial stroke severity (National Institutes of Health Score, NIHSS on admission), hemorrhagic transformation on follow-up imaging after one day, and clinical outcome (mRS and NIHSS after 3 months) (Fig. 6e). Patients with early venous filling had significantly lower collateral scores (p= 0.002) and more remaining neurological deficits at three months (NIHSS 1 (IQR4) versus 0 (IQR 0), p= 0.041) compared to those without early venous filling. Furthermore, hemorrhagic transformation of the infarcted area was detected only in patients with early venous filling (47.8% versus 0%; p = 0.007), suggesting a reduction in structural integrity of the vasculature in the infarcted areas.

**Fig. 6:**
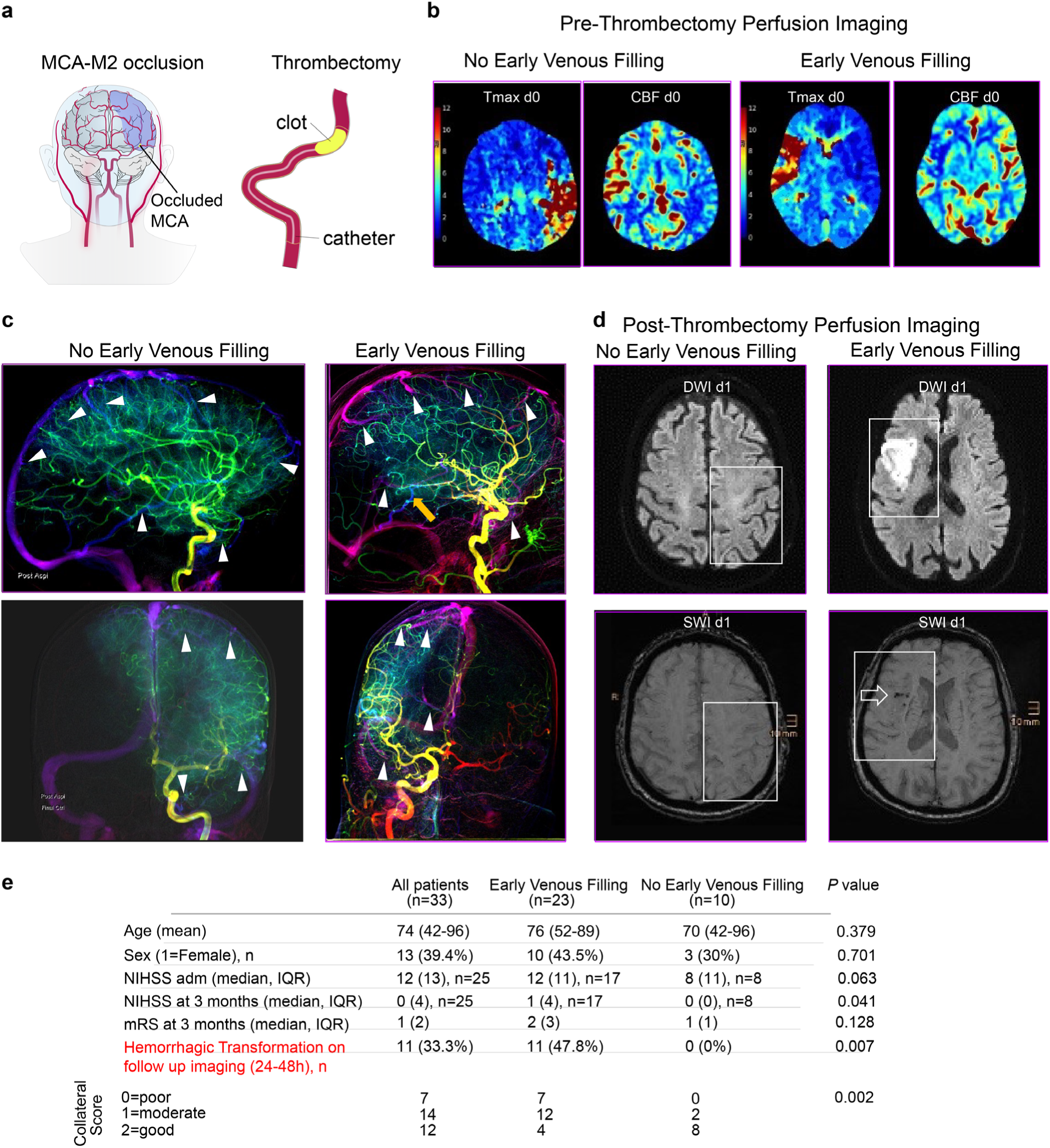
Rapid reperfusion after thrombectomy in stroke patients a) Schematic representation of the location of MCA-M2 occlusion in stroke patients. b) Pre-thrombectomy CT perfusion imaging in two stroke patients with distal MCA-M1 occlusion. Both have similar extent and severity of the perfusion deficit on Tmax maps (a Tmax > 6 sec corresponds to the ischemic core area), and corresponding CBF maps. Left patient (MCA-M2 occlusion on the left) had no early venous filling post thrombectomy, right patient MCA-M2 occlusion the right) showed early venous filling after thrombectomy. c) Color coded MIP (maximum intensity projection) DSA data from a patient without (left) and with (right) early venous filling indicated using white arrowheads. d) MRI 1 day after thrombectomy from the same two patients as in b) shows early infarct extension on diffusion-weighted images (DWI; white area) within the former hypoperfused area (white box). Below: corresponding slice from Susceptibility Weighted Image (SWI), where blood deposits appear black (arrow on the right side). Note that the patient with early venous filling has a considerably larger infarct on DWI and hemorrhagic transformation on SWI.

## Discussion

Futile recanalization, i.e. insufficient clinical and tissue recovery despite successful clot removal in stroke, is an immense clinical problem. Its underlying mechanisms remain poorly understood. Insufficient reperfusion of the microvascular bed or accelerated tissue injury upon reperfusion have been previously discussed ^25–27^. Here, we assessed tissue perfusion and arteriole reactivity during stroke and recanalization treatment in three mouse strains with genetic background-dependent differences in the number of native collaterals/LMCs interconnecting their MCA and ACA territories. We show that upon stroke induction, different levels of blood flow were recruited across the LMCs, depending on the number of collaterals present (i.e., “collateral status”) to the ischemic territory. Infarct volume was smaller and stroke-induced functional deficits were less severe in mice with abundant LMCs. Of particular importance, we found that upon thrombolysis, poor collateral status was associated with a fast and overshooting reperfusion, culminating in more severe tissue injury and worse clinical deficit. Data from the mouse model and from stroke patients suggest that abundant LMCs are involved in cushioning this potentially harmful reperfusion response.

Previous studies employing other stroke models suggested that collaterals act as alternative routes for blood flow supply ^28, 29^. Our two-photon microscopy findings add to this by demonstrating collateral flow and diameter increases directly after stroke, as well as a response to reperfusion treatments. Since most proximal arterial branches are not assessable by conventional imaging techniques, we used uUS (Fig.3d-f), which can provide hemodynamic imaging of the whole brain ^30, 31^. By combining ultrasound with two-photon microscopy and LSCI, we established a unique picture of the dynamic changes in diameter and flow in both the collaterals themselves as well as the vessel segments in the distal MCA that they perfuse during occlusion, as well as collateral flow, vascular integrity, and reperfusion patterns from the single-vessel level to deep brain areas.

In the thrombin model of stroke and thrombolysis, we observed significant differences in tissue reperfusion after recanalization. In mice with poor LMC networks, a rapid and steep CBF increase upon recanalization caused hyperperfusion of ischemic areas, hemorrhagic complications, and unfavorable outcomes. The overshoot in CBF above baseline levels observed during reperfusion in deeper, M3-supplied areas most likely resembles a loss of regulatory capacity in the affected vessels. Impaired vasodilatory capacity in response to a CO_2_ stimulus in mice with poor or no LMCs confirmed this hypothesis. Thus, uncontrolled reperfusion may endanger the previously ischemic brain tissue and induce reperfusion injury.

Reperfusion injury is a recognized problem in ischemic heart disease ^32^. Several mechanisms have been implicated in mediating excess myocardial injury in response to coronary artery recanalization, among them mitochondrial dysfunction with the formation of reactive oxygen species, capillary and myocyte swelling, microvascular leucocyte adhesion and no reflow, as well as and increased capillary permeability ^33–35^. Although ischemia in the heart and brain follow common grounds ^36^, there has been no description of altered brain perfusion potentially resembling reperfusion injury in rodent models, and no convincing evidence of reperfusion injury in stroke patients, so that the clinical validity of this concept was recently questioned ^37^. We here provide evidence of a potentially harmful hyperemic reperfusion in stroke based on i) dynamic flow measurements capturing hyperperfusion, ii) decomposition of vascular structures after recanalization (constriction of M5-branches on 2-photon microscopy), and iii) diminished vascular function (hypercapnia challenge) in mice with poor LMCs.

We cannot rule out that strain-specific factors other than the extent of LMC could explain differences in reperfusion patterns between Balb-C mice and the other two strains (e.g., differences in coagulation factors or individual resistance to ischemia). However, Rabep2-/- mice shared the exact same genetic background compared to C57Bl/6 mice except for the mutant gene Rabep2, arguing against other variables significantly affecting the observed difference in reperfusion response. This is further supported by data obtained from the Rabep2+/+ littermates. In our model, recanalization success was independent of LMC status. The thrombin model is a clinical relevant model of stroke, clot formation and lysis as well as vascular responses to intravenous thrombolysis, using the same rt-PA dose and infusion rate as in patients. Other animal models of stroke using filaments, ligatures, clamping, or electrocoagulation do not capture the processes involved in the recanalization of a clot and thrombolysis-induced reperfusion ^38–40^. While extent of collaterals was previously shown to correlate with thrombus length^41^ and fibrinolysis resistance ^42^, our model with very fresh and homogenous clots may not reflect heterogeneity in clot composition and source present in stroke patients. However, thus we were able to observe reperfusion dynamics in relation to collateral status independent of clot composition.

Strikingly, we demonstrated a similar rapid reperfusion pattern in patients with poor collaterals and ischemic stroke who had undergone recanalization through thrombectomy. We used color-coding according to DSA contrast arrival times directly post-thrombectomy in patients with stroke due to distal MCA-M1 or MCA-M2 occlusion. Early venous filling, albeit investigated with a different analysis approach and in a more heterogeneous stroke patient cohort, was previously suggested to indicate harmful hyperperfusion following thrombectomy ^43^. In our patient cohort, this was reflected by more severe neurological deficit (NIHSS), higher disability (mRS), and hemorrhagic transformation of the infarcted area on follow-up imaging. Hemorrhages on SWI-MRI are indicators of blood-brain-barrier disruption, one of the hallmarks of secondary infarct complications ^44^. Therefore, early venous filling may indicate patients at risk of ICH and futile recanalization immediately post-thrombectomy, thus calling for intensive post-interventional medical monitoring.

Our findings have important implications for stroke treatment. LMCs are crucial for preventing reperfusion injury arising from recanalization. We show that patients with poor LMCs are more likely to suffer from severe stroke due to a lack of compensatory intra-ischemic perfusion. Importantly, we demonstrate that patients with poor LMCs are at particular risk of rapid reperfusion associated with more severe tissue damage and hemorrhagic infarct transformation after stroke treatments. LMC status should be considered as an indicator of vulnerable stroke patients in stroke trials. Longer times to treatment could be exceedingly dangerous in these patients. Vascular risk factors are likely to affect LMC function, which should be investigated further. In addition to their potential use as biomarkers for outcome of stroke treatments, LMCs may be targeted for stroke treatments. In the IMPACT 24 trial, flow increase in ipsilateral collaterals induced by sphenopalatine ganglion stimulation showed promising results for stroke patients with cortical stroke in the anterior circulation ^45^. In addition to fostering LMC function, another feasible approach would be slowing down rapid reperfusion (e.g., by graded or staccato removal of the obstructive clot in patients with poor LMCs), thus reducing tissue injury and improving stroke patient outcome.

## Online Methods

### Animals

For all experiments, Balb-C (Charles Rivers, no. 028), C57BL/6 (Charles Rivers, no. 000664) and Rabep2-/- ^16^ mice, mixed female/male, between 3 to 4 months of age were used. The animals had free access to water and food and an inverted 12-hour light/dark cycle to perform experiments during the dark, active phase. All experiments were approved by the local veterinary authorities in Zurich and conformed to the guidelines of the Swiss Animal Protection Law, Veterinary Office, Canton of Zurich (Act of Animal Protection 16 December 2005 and Animal Protection Ordinance 23 April 2008), animal welfare assurance numbers ZH165/19 and ZH224/15.

### Anaesthesia

For head-post and cranial window implantation, mice were anesthetized intraperitoneally with a mixture of fentanyl (0.05 mg/kg bodyweight; Sintenyl, Sintetica), midazolam (5 mg/kg bodyweight; Dormicum, Roche), and medetomidine (0.5 mg/kg bodyweight; Domitor, Orion Pharma). Throughout anaesthesia, a facemask provided oxygen at a rate of 500 ml/min. For stroke surgery and two-photon imaging anaesthesia was induced with isoflurane 4%, maintained at 1.2% with continuous supply of an oxygen/air mixture (30%/70%). For LSCI and ultrafast ultrasound recordings, anaesthesia was changed to a mixture of fentanyl (0.05 mg/kg bodyweight; Sintenyl, Sintetica), midazolam (5 mg/kg bodyweight; Dormicum, Roche), and medetomidine (0.5 mg/kg bodyweight; Domitor, Orion Pharma) before stroke induction. The core temperature of the animals were kept constant at 37°C using a homeothermic blanket heating system during all surgical and experimental procedures (Harvard Apparatus). The animal’s head was fixed in a stereotaxic apparatus, and the eyes were kept moist with a vitamin A eye cream (Bausch & Lomb).

### Head-Post Implantation

A bonding agent (Gluma Comfort Bond; Heraeus Kulzer) was applied to the cleaned skull and polymerized with a handheld blue light source (600 mW/cm^2^; Demetron LC). A custom-made aluminum head post was connected to the bonding agent with dental cement (EvoFlow; Ivoclar Vivadent AG) for stable and reproducible fixation in the microscope setup. The skin lesion was treated with an antibiotic cream (Fucidin®, LEO Pharma GmbH)) and closed with acrylic glue (Histoacryl, B. Braun). After surgery, animals were kept warm and received analgesics (buprenorphine 0.1 mg/kg bodyweight; Sintetica).

### Cranial window surgery

A 4 x 4 mm craniotomy was performed above the somatosensory cortex (centered above the left somatosensory cortex) with a dental drill (Bien-Air). A square coverslip (3 x 3 mm, UQG Optics) was placed on the exposed dura mater and fixed to the skull with dental cement ^46^.

### Thrombin stroke model

We induced a focal cerebral ischemia using the thrombin-model of stroke. The procedure steps for the cerebral ischemia induction were performed as described previously ^20, 21, 47^. In brief, mice heads were fixed in a stereotactic frame, the skin between the left eye and ear was incised and the left temporal muscle retracted. After craniotomy above the middle cerebral artery (MCA) M2 segment, the dura was removed, and a glass pipette (calibrated at 15 mm/μl; Assistant ref. 555/5; Hoechst, Sondheim-Rhoen, Germany) was introduced into the lumen of the MCA. Through the pipette, 1μl of purified human alpha-thrombin (1UI; HCT-0020, Haematologic Technologies Inc., USA) was injected to induce the formation of a clot in situ. Ten minutes after thrombin injection, the pipette was removed. Ischemia induction was considered successful when CBF rapidly dropped to at least 50 % of baseline level in the MCA territory ^20, 48^ and remained below 50% for at least 30 minutes. Animals without stable ischemia induction were excluded from further experiments.

### Rt-PA administration

Thirty minutes after ischemia induction, thrombolysis was initiated via tail vein injection of rt- PA (10 mg/kg, Actilyse, Boehringer Ingelheim). According to previous data ^20, 49^, ten percent were given as a bolus and ninety percent were perfused at a body weight dependent perfusion rate between 6 and 9.3µl/min. Control groups received the vehicle (0.9% saline) instead of rt- PA.

### Laser Speckle Cortical Imaging (LSCI)

Cortical relative cerebral blood flow (CBF) changes were monitored before and during ischemia, and throughout the recanalization phase for 110 min using a laser speckle contrast imaging monitor (FLPI, Moor Instruments, UK). The acquisition was performed with a frame rate of 0.25 Hz. Afterwards the LSCI images were exported and generated with arbitrary units in a 256-colour palette by the MOOR-FLPI software.

### Two-photon imaging

After cranial window implantation, mice could recover for 1-2 weeks prior to two-photon imaging. Imaging was performed using a custom-built two-photon laser scanning microscope (2PLSM) ^50^ with a tuneable pulsed laser (Chameleon Discovery TPC, Coherent Inc.) equipped with a 20x (W-Plan-Apochromat 20x/1.0 NA, Zeiss) water immersion objective. During measurements, the animals were head-fixed and kept under anaesthesia as described above. Vasculature was labelled with intravenous injection of FITC–dextran (2-MDa, Sigma-Aldrich; FD2000S) or Texas red Dextran (70 MDa, Life Technologies catalogue number D-1864), ten minutes before imaging. To visualize arterioles specifically, Hydrazide (Alexa Fluor™ 633 Hydrazide Life Technologies catalogue number A30634) was injected intravenously. Emission was detected with GaAsP photomultiplier modules (Hamamatsu Photonics) equipped with 475/64, 520/50 nm and 607/70 band pass filters and separated by506, 560 and 652 nm dichroic mirrors (BrightLine; Semrock). The microscope was controlled by a customized version of ScanImage (r3.8.1; Janelia Research Campus ^51^). Baseline measurements were performed 90 min before induction of ischemia. Different branches from the MCA were identified based on the vascular anatomy. For acquisition of red blood cell (RBC) velocity and vessel diameters, line scans were performed in arterioles at 11.84 Hz, 0.55 μm/pixel and for 12.7 s. Three regions of interest (ROIs) were identified on the terminal distal branches of arterioles from the MCA. Three other ROIs were placed on collaterals, identified (1) as the arterio-arterial anastomosis connecting MCA to ACA and (2) by their typical tortuosity and low flow. The same vessels were evaluated before stroke, after stroke and throughout the experiment.

Line scans were processed with a custom-designed image processing toolbox for MATLAB (Cellular and Hemodynamic Image Processing Suite; R2014b; doi: 10.1007/s12021-017-9344-y; MathWorks). Vessel diameters were determined at full width half maximum (FWHM) from a Gaussian fitted intensity profile drawn perpendicular to the vessel axis. RBC velocity flow was calculated with the Radon algorithm.

### Ultrafast ultrasound imaging

Ultrafast ultrasound imaging (uUS) was performed on a prototype research ultrafast scanner with Neuroscan live acquisition software (ART Inserm U1273, Paris, France & Iconeus, Paris, France) using a 15 MHz ultrasound probe (0.11 mm pitch) which enables a 110 μm x 100 μm in plane resolution with a depth of 10 mm. The probe was mounted on 4 motors (3 translation + 1 rotation) for automatic positioning and scans. During measurements, the animals were head-fixed and kept under anaesthesia as described above.

### 4D Ultrasensitive Doppler

For each ultrafast Doppler image, 200 compounded frames (11 angles between −10°: 10°) were acquired at 500 Hz. Singular Value Decomposition clutter filters were used and the 60 first singular values were removed to separate blood signal from tissues. The energy in each pixel was calculated to form a Power Doppler image. The motor was then moved to cover 24 coronal slices imaged every 0.3 mm to reconstruct a 10 mm x 12.8 mm x 8 mm volume between β+2mm and β-6mm with a temporal resolution of 40 seconds. We performed a 10 minute recording for the baseline and two sets of recordings of 60 minutes each after stroke induction.

### Automatic probe positioning

To enhance the reproducibility from one acquisition to another, the probe (before starting each 4D acquisition) was automatically positioned with the Brain Positioning System software (Nouhoum et al. 2021) (Iconeus, Paris, France) over a coronal slice at Bregma minus 2.3 mm and aligned automatically to the Allen brain atlas.

### Data processing

Within each acquisition (baseline, post-stroke1, post-stroke2), the volumes were realigned on the temporal average volume. Then, for each set of three acquisitions, a merged acquisition was created after registration of post-stroke1 and post-stroke2 acquisitions on the baseline and concatenation of baseline and the resulting registered post-stroke1 and post-stroke2 acquisitions.

### Vascular Territories (VT) segmentation and CBV time-profile extraction

VT segmentation was performed manually on a baseline angiography reference ^52^. Then, all the merged acquisitions were registered on this vascular reference. Hence, the same VT segmentation was used for all animals. Mean CBV time-profiles inside each Region of Interest (ROI) were extracted and normalized by the temporal average of the baseline acquisition to obtain relative-CBV (rCBV) curves.

### Ultrasound Localization Microscopy

For each ultrasound localization microscopy (ULM) image, 100 μL of Sonovue® microbubbles (SonoVue, Bracco, Italy) were injected through a tail vein catheter. Every second 180 blocks of 800 compounded frames (3 angles at −5° 0° 5°) at 1000 Hz were acquired and reconstructed by a dedicated software (ART Inserm U1273, Paris, France & Iconeus, Paris, France). Briefly, a Singular Value Decomposition filter (removal of the 10 first singular values) was used to separate microbubble echoes from tissues and microbubble centroid positions were localized^30^. Microbubbles were tracked through consecutive frames, a tracking algorithm based on the Hungarian method for assignment. Tracks were interpolated and a density image was reconstructed by counting the number of tracks in each pixel of the grid. ULM was performed for 180 seconds before as well as one hour and two hours after stroke induction.

### CO_2_ challenge

To evoke hypercapnic hyperemia, mice were subjected to inhalation of 7.5% CO_2_ (PanGas) for five minutes while anaesthetized with triple anaesthesia. At the same time, to assess cerebrovascular reactivity, blood flow was monitored using LSCI. The CO_2_ challenge was administered once before induction of stroke and 60 minutes post-stroke. The maximum dilation of an artery was measured by evaluating an ROI on the most distal MCA segment (MCA-M5 supplied territory).

### Tissue clearing

Mouse brains were stained for arteries and cleared using a modified version of the iDISCO protocol ^18^. Mice were anesthetized, perfused, and the brains post-fixed in 4% PFA in PBS for 4.5 hours at 4°C, shaking at 40 rpm. Mouse brains were washed in PBS for 3 days at RT and 40 rpm, with daily solution exchange. Samples were dehydrated in serial incubations of 20%, 40%, 60%, 80% methanol (MeOH) in ddH_2_O, followed by 2 times 100% MeOH, each for 1 hour at RT and 40 rpm. Pre-clearing was performed in 33% MeOH in dichloromethane (DCM) overnight (o.n.) at RT and 40 rpm. After 2 times washing in 100% MeOH each for 1 hour at RT and then 4°C at 40 rpm, bleaching was performed in 5% hydrogen peroxide in MeOH for 20 hours at 4°C and 40 rpm. Samples were rehydrated in serial incubations of 80%, 60%, 40%, and 20% MeOH in in ddH_2_O, followed by PBS, each for 1 hour at RT and 40 rpm. Permeabilization was performed by incubating the mouse brains 2 times in 0.2% TritonX-100 in PBS each for 1 hour at RT and 40 rpm, followed by incubation in 0.2% TritonX-100 + 10% dimethyl sulfoxide (DMSO) + 2.3% glycine + 0.1% sodium azide (NaN3) in PBS for 5 days at 37°C and 65 rpm. Blocking was performed in 0.2% Tween-20 + 0.1% heparin (10 mg/ml) + 5% DMSO + 6% donkey serum in PBS for 2 days at 37°C and 65 rpm. Samples were stained gradually with Cy3-conjugated monoclonal anti-α-smooth muscle actin antibody (Sigma Aldrich, C6198) 1:800 in 0.2% Tween-20 + 0.1% heparin + 5% DMSO + 0.1% NaN3 in PBS (staining buffer) in a total volume of 1.5 ml per sample every week for 4 weeks at 37°C and 65 rpm. Washing steps were performed in staining buffer 5 times each for 1 hour, and then for 2 days at RT and 40 rpm. Clearing was started by dehydrating the samples in serial MeOH incubations as described above. Delipidation was performed in 33% MeOH in DCM o.n. at RT and 40 rpm, followed by 2 times 100% DCM each for 30 minutes at RT and 40 rpm. Refractive index (RI) matching was achieved in dibenzyl ether (DBE, RI = 1.56) for 4 hours at RT.

### Whole-brain imaging

Three-D stacks of cleared brains were acquired using a custom-made selective plane illumination microscope (meso-SPIM, version V5) ^53^ (http://www.mesospim.org). One x zooms with a field of view of 1.3 cm and isotropic resolution of 6 µm/voxel. Imaging data were post-processed using Fiji (Image J, 1.8.0_172 64 bit) and Imaris (Oxford Instruments, 9.8.0).

### Quantification of lesion volume and hemorrhages

At day 7 post stroke, mice were euthanized by receiving an overdose i.p. injection of pentobarbital (200 mg/kg) followed by decapitation. Brains were extracted and cut into 1 mm thick coronal slices from 6.5 to 0.5 mm anterior to the inter-aural line and placed in 2% 2,3,5-triphenyltetrazolium chloride (TTC, cat. #T8877, Sigma-Aldrich, St. Louis, MO) for 10 min at 37 °C. Infarct areas were determined by a blinded investigator using an image analysis system (Image J version 1.41). To correct for brain swelling, each infarct area was multiplied by the ratio of the surface of the intact (contralateral) hemisphere to the infarcted (ipsilateral) hemisphere at the same level. Total volume of damaged tissue, expressed as cubic millimetres, was calculated by linear integration of the corrected lesion areas ^54^. The presence of hemorrhage was recorded from TTC-stained brain slices at the time of premature death or sacrifice at day 7. Microscopic, macroscopic and total hemorrhagic scores were visually quantified on each level, from TTC-stained brain slices, as previously described ^55^.

### Behavioral assessment

Before stroke as well as on day 1, 3 and 7 post ischemia, neurological deficits were assessed using the adhesive tape removal test and a composite observational neurological score. In the adhesive tape removal test an investigator blinded to treatment applied two rectangular tape strips (0.3 × 0.4 cm) to both forepaws. In order to assess sensorimotor deficits, time was recorded until the animals first had contact (sensory function/neglect) and removed (motor function) both stripes ^56^. Before stroke, mice were trained to remove both tapes within 10 seconds. In addition to the sticky tape test, the neurological score was obtained using a composite grading score as described before ^20^. A lower score indicates larger neurological deficits, while a score of 13 points indicates no neurological deficit.

### Patient cohort and clinical parameters

Patient data analysis was performed according to the ethical guidelines of the Canton of Zurich with approval from the local ethics committee (“PREDICT”; KEK-ZH-Nr. 2014-0304). From the Swiss Stroke Registry, we screened patients treated with acute ischemic stroke at the Department of Neurology, University Hospital Zurich from 01-04 2021 (n=202). Treatment decisions including thrombectomy were made by treating physicians according to current clinical guidelines. Stroke imaging including CT angiography (CTA), DSA and CT perfusion (CTP) was performed according to routine clinical protocols. We selected only those patients with distal MCA-M1 or M2 occlusion (one open MCA branch had to be identified for reference), who were sent to thrombectomy and had a successful recanalization result, corresponding to mTICI (modified treatment in cerebral infarction) grade 2b/3 (n= 26) ^26^. We excluded patients who objected to the use of their data for research purposes, or patients without any clinical follow-up information. Five patients were excluded because of insufficient quality of DSA data for semi-quantitative analysis, so that 21 patients were included.

Clinical information about sex, and age, and information about stroke severity according to the National Institute of Health Stroke Scale (NIHSS)^57^ on admission as well as follow up NIHSS and disability according to the modified Rankin Scale (mRS)^58^ at 3 months post stroke were derived from the registry. Information about hemorrhagic transformation within the ischemic area was taken from clinical neuroradiology reports of brain MRI acquired one day after the intervention. Classification of LMC status as poor, intermediate, or good was done according to EXTEND-IA criteria ^59^.

### DSA data analysis

We collected biplane angiography runs obtained during the thrombectomy procedure with the in house angio-suite (Axiom-ArtisQ and Axiom-Artis Zee, biplane, Siemens, Germany) from the included patients. The standard protocol for image acquisition was to divide fluoroscopic acquisition into 3 phases; 10 images with a frame rate of 500ms, 5 images with 1s frame rate, and the remaining images with a frame rate of 2s for dose reduction. From each patient, only the image series from the final control runs where utilized. The corresponding DICOM images where anonymized and transferred to Fiji (Image J, 1.8.0_172 64 bit) for semiquantative analysis. Rather than specifying “arterial” or “parenchymal” early venous filling ^43^, we used the open source “time lapse color coder” module from ImageJ to allow visual representation of contrast filling delays in angiographic images. To do this, the first frame of arterial filling till the last frame of the DSA images were manually selected and each frame was coloured according to a pseudocolor LUT (look up table) to display even small time delays with maximum visibility. As all our cases had at least one non-occluded MCA branch, we took this territory as an internal reference for venous delay in this patient. Dichotomous visual assessment of venous filling delays into “early venous filling” (i.e., clear difference of one or more venous drainage territories of the affected hemisphere) and “no early venous filling” was done on maximum intensity projections of the color-coded time series based on apparent temporal encoded colour differences in venous delay. Two interventional neuroradiologists, unaware of clinical or imaging outcome, performed the readings and assessments independently.

### Statistical analysis

Data in all groups was tested for normality using D’Agostino-Pearson omnibus normality test. Parametric statistics were used only if the data in all groups in the comparison were normally distributed. Statistical analysis was performed using the GraphPad Prism (version 8.0; GraphPad Software La Jolla, CA, USA). All statistical tests and group size (n) are indicated in the figures. To account for multiple observations within the imaged mouse, results from each group were compared using univariate nested model t-tests before proceeding with the discriminant analysis. Results were expressed either as mean ± s.e.m. (standard error of mean), or median (interquartile range) in box plots. Comparisons between patient groups were done using IBM SPSS Statistics (Version 27, 28). Significance (P < 0.05) between two groups was calculated using unpaired Student’s t-test or paired t-test for normally distributed data, or with the Mann–Whitney test for data with non-normal distribution. Frequencies were compared using the Chi Square test.

## Acknowledgements

We like to thank James Faber, University of North Caroline Chapel Hill, for his conceptual advice and review of the manuscript.

## Funding

The Swiss National Science Foundation (PP00P3_170683 and PP00P3_202663), the Swiss Heart Foundation, the UZH Clinical Research Priority Program stroke, the Betty and David Koetser Foundation, and a ZNZ PhD grant funded this work.

## Supplementary Figures

**Supplementary Fig. 1:**
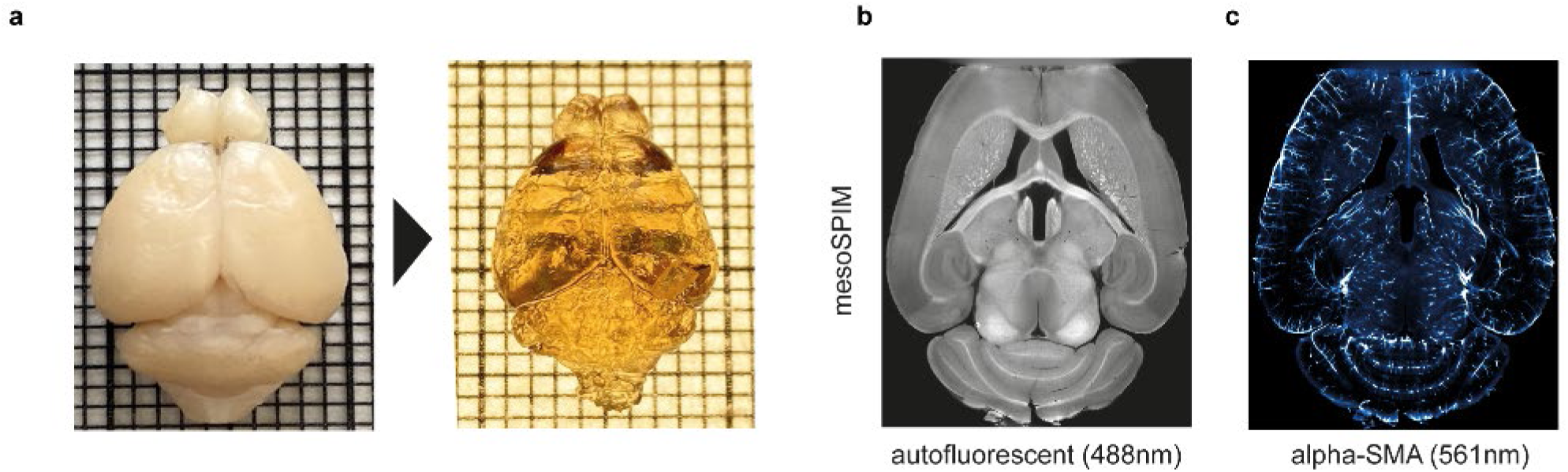
IDISCO clearing protocol (**a**) PFA perfused (left) and iDISCO cleared (right) mouse brain; Representative image of autofluorescence (**b**) and alpha-SMA signal (**c**) of iDISCO cleared and anti-SMA-Cy3 stained brain using light sheet microscopy.

**Supplementary Fig. 2:**
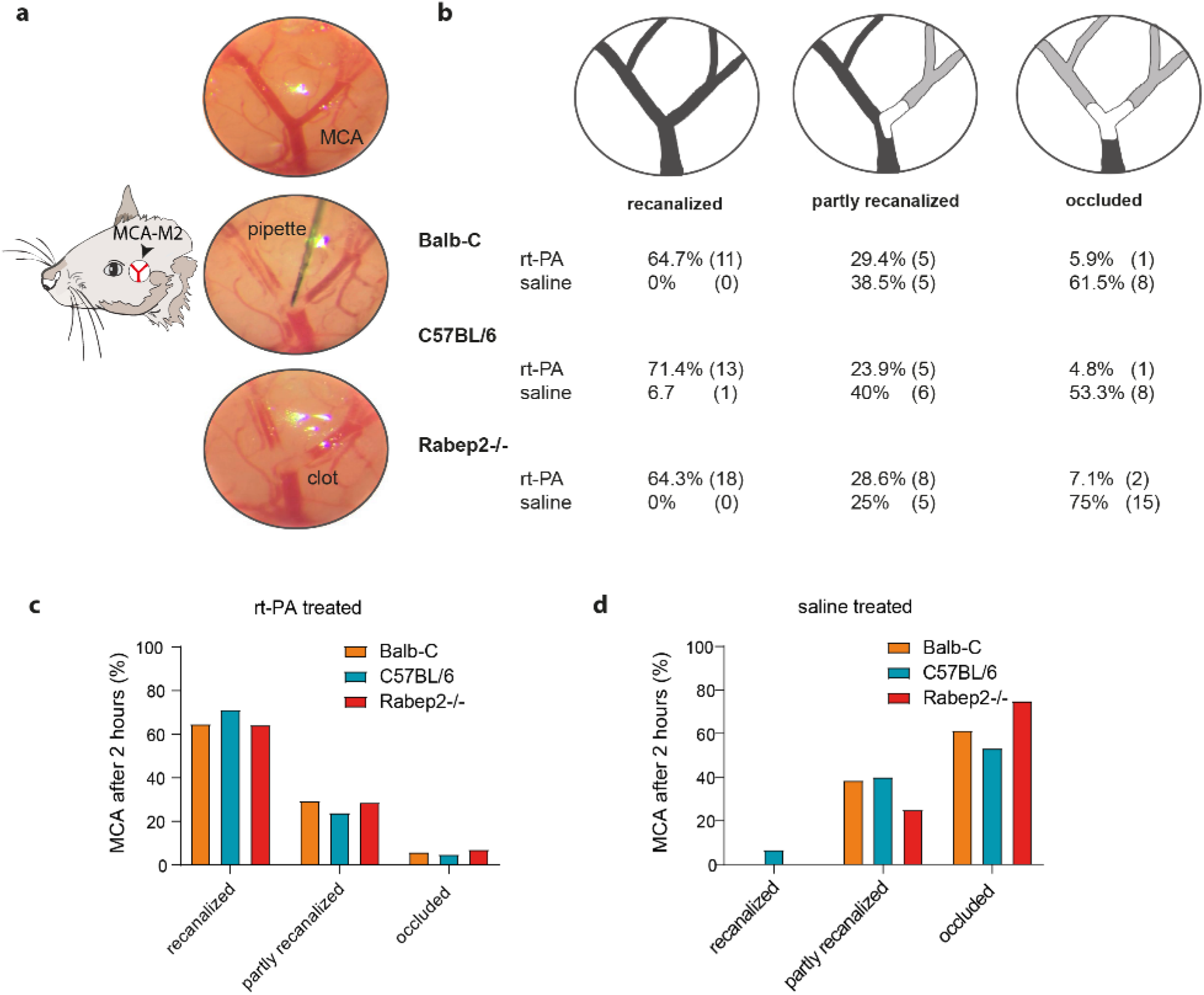
Thrombin model of stroke and recanalization rates upon rt-PA or control (saline) treatment. (**a**) Thrombin model of stroke; clot formation through thrombin injection into the MCA-M2 bifurcation visualized with 45x zoom microscope. (**b**) Schematic drawing of the three recanalization conditions of the MCA-M2 bifurcation two hours post stroke. The table displays the results of MCA status in either rt-PA treated animals or saline treated controls. N Numbers are displayed in brackets. (**c**)- (**d**) The graphs show strain differences in rt-PA (**c**) or saline (**d**) treated animals. Recanalization rates do not differ between Balb-C, C57BL/6 and Rabep2-/-; p= 0.468 for saline treatment and p = 0.998 for rt-PA treatment, two sided significance in Pearson Chi square test.

**Supplementary Fig. 3:**
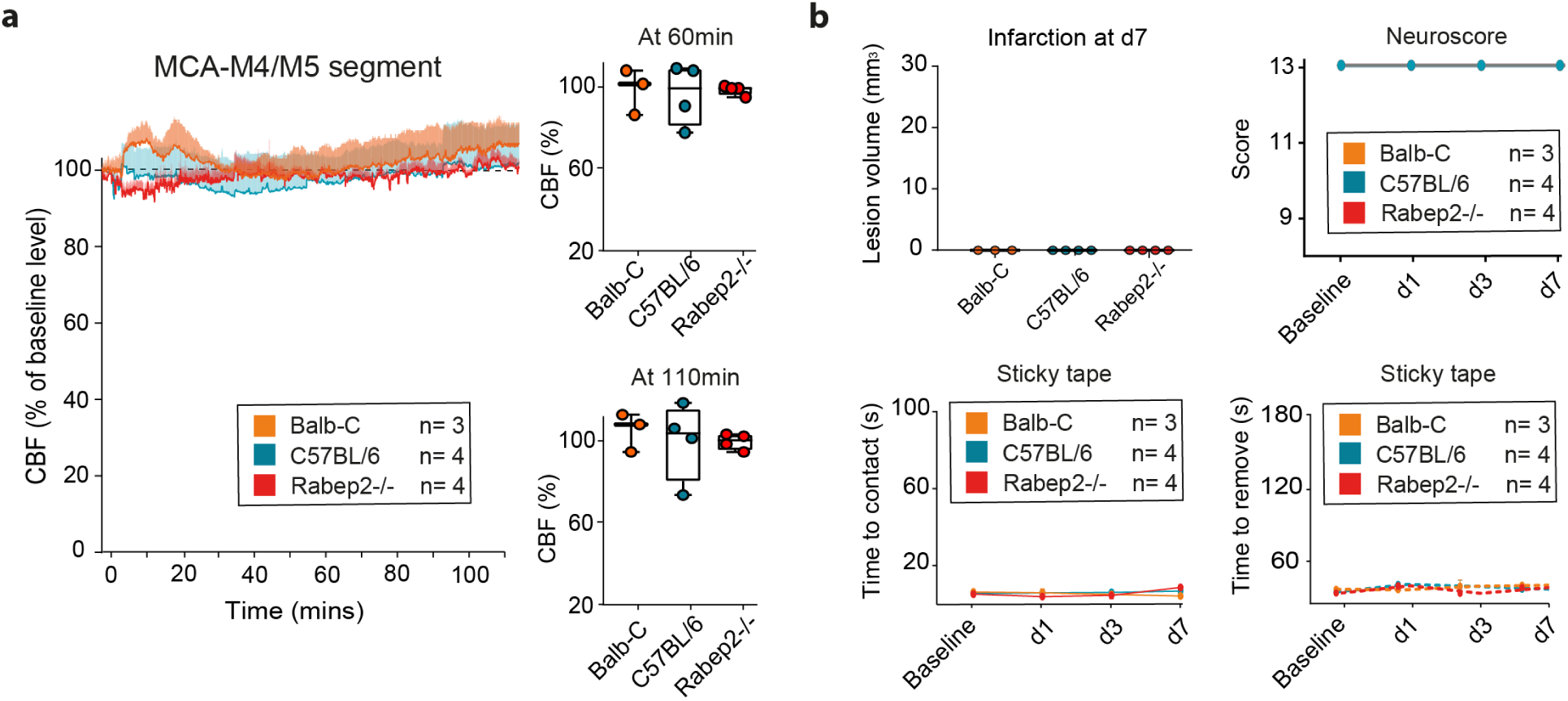
Sham operated animals displaying no CBF changes on LSCI analysis or neurological deficits. (**a**) LSCI analysis of MCA-M4/M5 area of sham operated Balb-C, C57BL/6 and Rabep2-/- mice; group sizes are indicated in the figure. (**b**) outcome including infarct volume on d7 as well as neurological assessment using neuroscore and sticky tape test.

**Supplementary Fig. 4:**
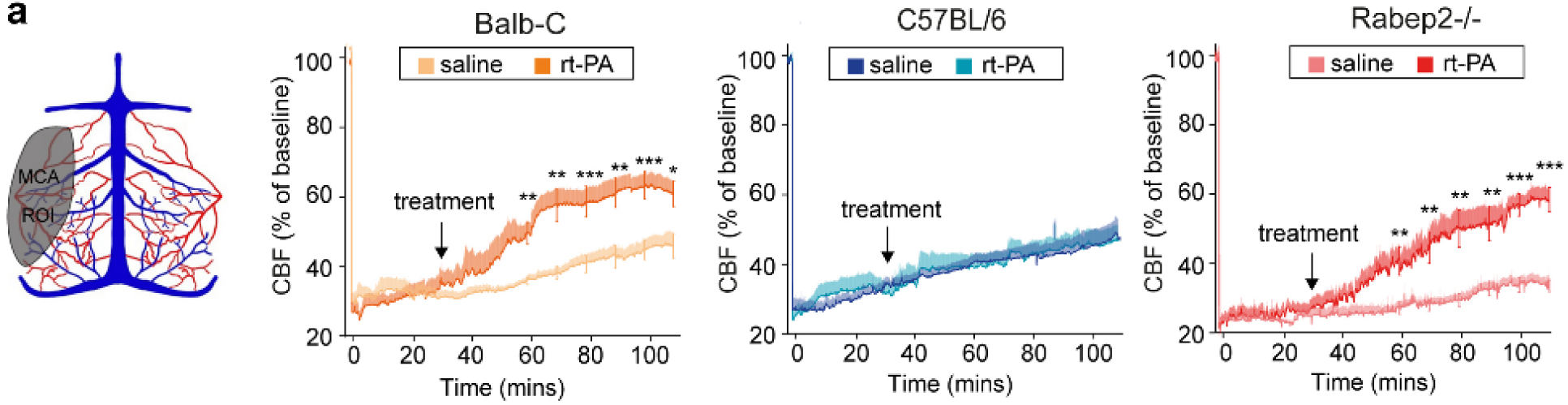
Reperfusion after stroke and thrombolysis assessed by LSCI. (**a**) Schematic drawing of the LSCI view on pial vessels and the MCA-M4/M5 region of interest (ROI) used for analyses. LSCI recordings for the MCA-M4/M5 ROI compared to baseline in saline or rt-PA treated Balb-C, C57BL/6, and Rabep2-/- mice, n = 10/group, ∗p < 0.05, ∗∗p < 0.01, ∗∗∗p < 0.001, two-tailed Mann-Whitney U-test.

**Supplementary Fig. 5:**
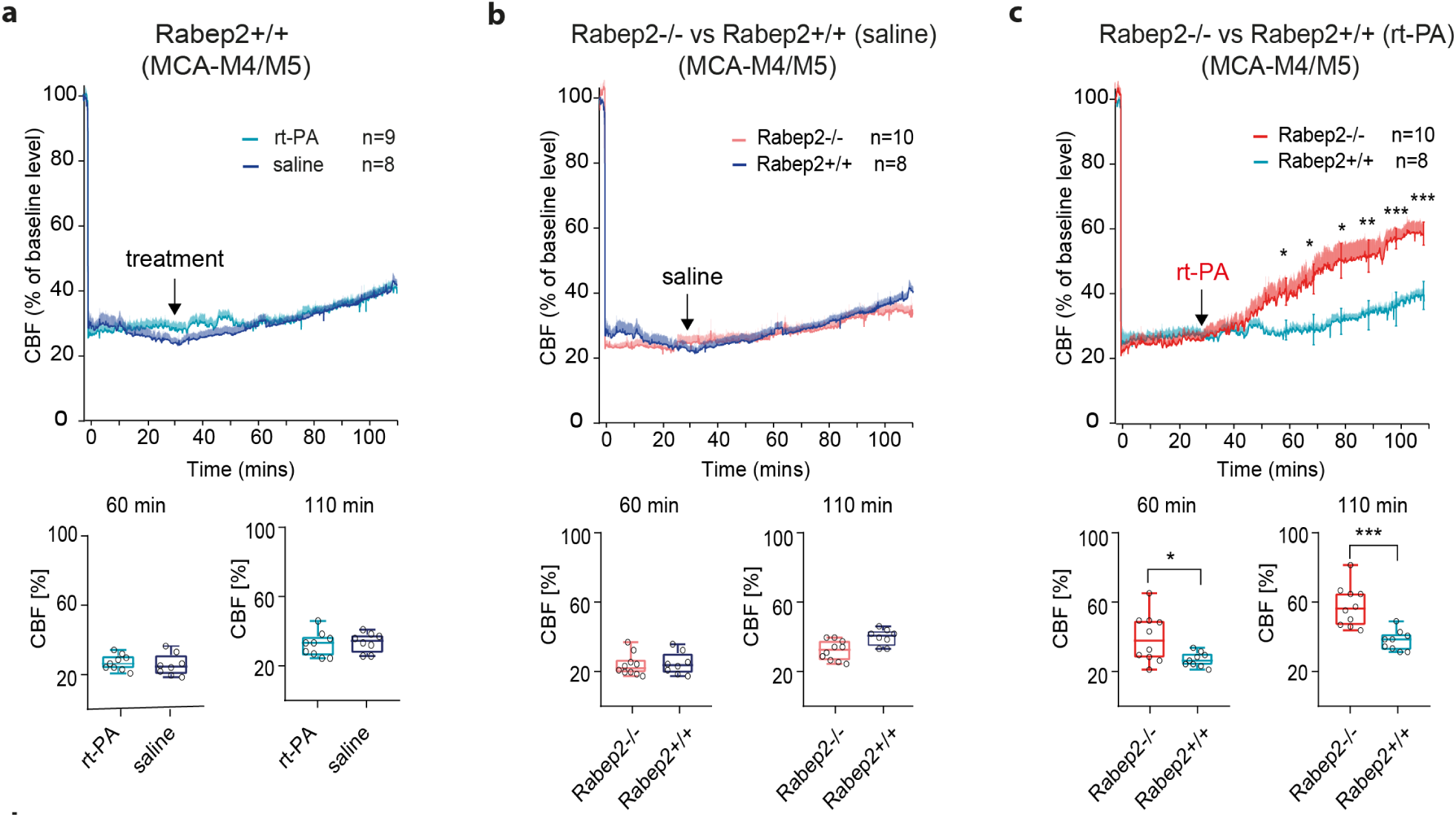
Rabep2+/+ analysis and comparison with Rabep2-/- mice; plus sex difference. **(a**) LSCI analysis for the MCA-M4 ROI compared to baseline in saline or rt-PA treated Rabep2+/+, in b) saline treated Rabep2+/+ and Rabep2-/- mice, and (**c**) rt-PA treated Rabep2+/+ and Rabep2-/- mice; group sizes are indicated in the figure, ∗p < 0.05, ∗∗p < 0.01, ∗∗∗p < 0.001, two-tailed Mann-Whitney U-test. (**d**) Bar graph depicting infarct volumes of individual mice at day 7, ∗p < 0.05, ∗∗∗p < 0.001 in two-tailed Mann-Whitney U-test.

**Supplementary Fig. 6:**
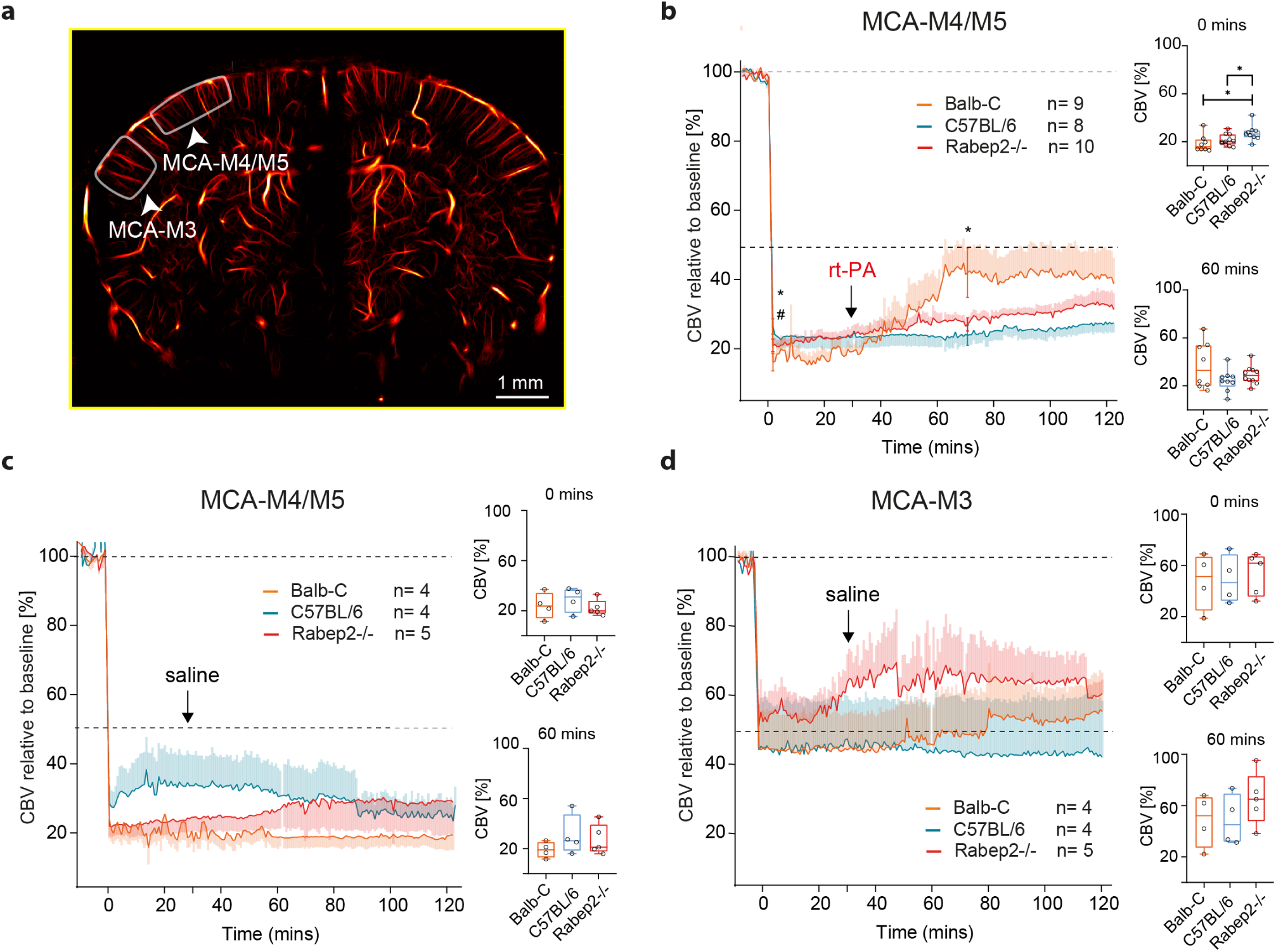
Ultrafast ultrasound (uUS) analysis of CBV in in the MCA-M3 and M4/M5 area in C57BL/6, Balb-C and Rabep2-/- mice. (**a**) ROI selection: MCA-M4/M5 and MCA-M3 territory in view of coronal section. 4D analysis (**b**) of the MCA-M4/M5 territory in rt-PA treated animals, (**c**) the MCA-M4/M5 territory and (**d**) the MCA-M3 territory of control treated animals; group sizes are indicated in the figure;∗p < 0.05, two-tailed Mann-Whitney U-test.

**Supplemental Fig. 7:**
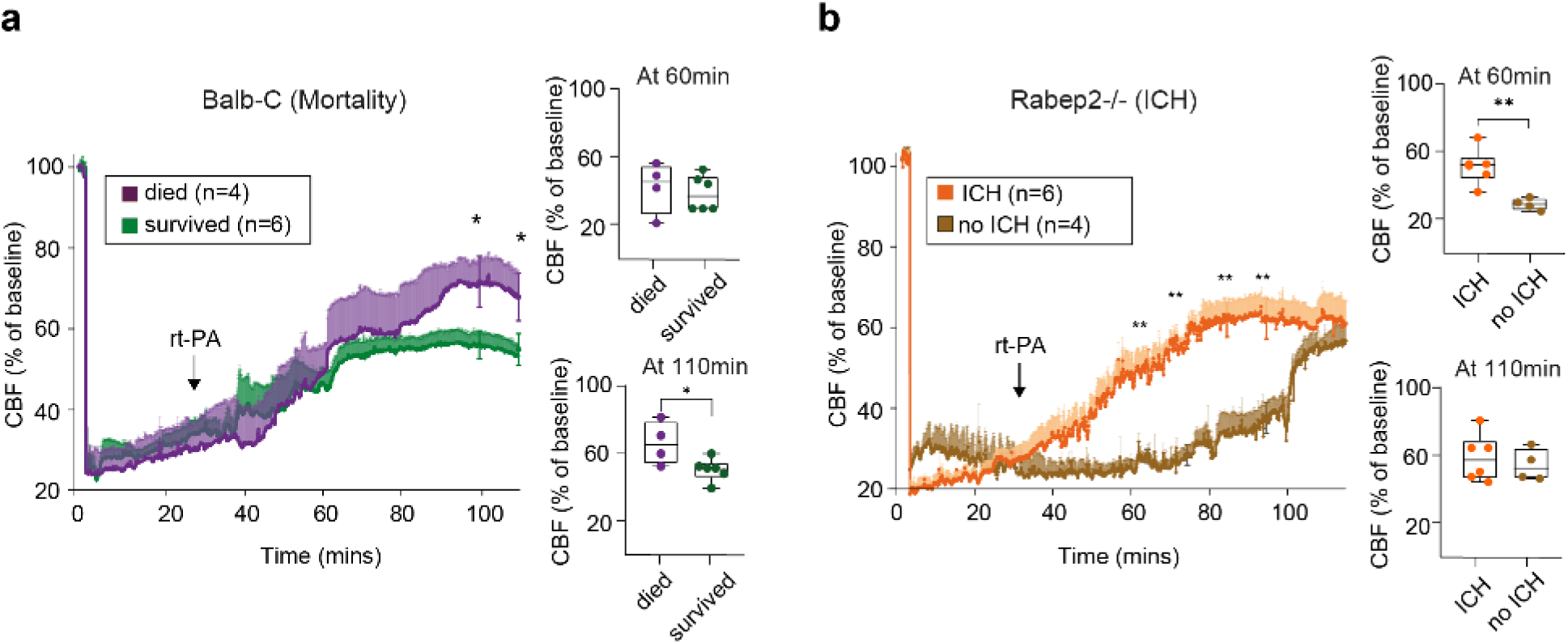
Complications after thrombolysis. Subgrouped analysis of LSCI data of the MCA-M4/M5 area from Balb-C and Rabep2 -/- mice (**a**) Subgrouped analysis of Balb-C mice, which died vs. survived treated with rt-PA; group sizes are indicated in the figure, ∗p < 0.05, two-tailed Mann-Whitney U-test. (**b**) analysis of Rabep2-/- mice displaying intracerebral hemorrhages (ICH) after thrombolysis versus those who did not, according to TTC; group sizes are indicated in the figure, ∗∗p < 0.01, two-tailed Mann-Whitney U-test.

**Supplemental Fig. 8:**
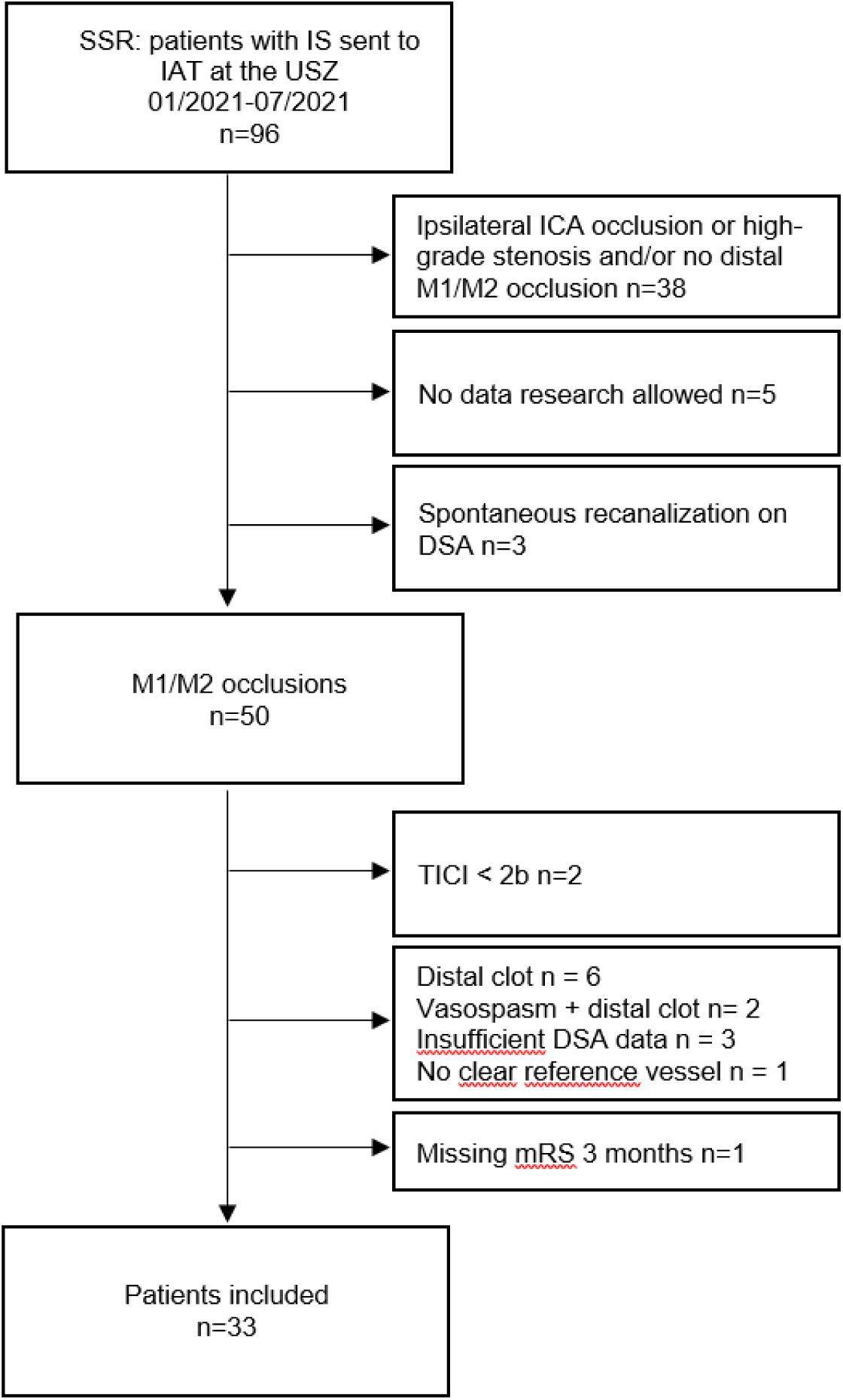
Patient Flow Chart Out of 96 patients treated with thrombectomy between 01- 07/2021, 33 patients with distal M1 or proximal M2 occlusions and sucessful recanalization (TICI >/= 2b-3) were included into the analysis. Reasons for exclusion are indicated. SSR: Swiss Stroke Registry. IS: Ischemic Stroke. IAT: intraarterial thrombectomy. USZ: University Hospital Zurich. ICA: Internal Carotid Artery.

## Notes

### Competing Interest Statement

The authors have declared no competing interest.

## References

1. Virani, S.S., et al. Heart Disease and Stroke Statistics-2021 Update: A Report From the American Heart Association. Circulation 143, e254–e743 (2021).

2. Campbell, B.C.V., et al. Ischaemic stroke. Nat Rev Dis Primers 5, 70 (2019).

3. Espinosa de Rueda, M., et al. Combined Multimodal Computed Tomography Score Correlates With Futile Recanalization After Thrombectomy in Patients With Acute Stroke. Stroke 46, 2517–2522 (2015).

4. Goyal, M., et al. Endovascular thrombectomy after large-vessel ischaemic stroke: a meta-analysis of individual patient data from five randomised trials. Lancet 387, 1723–1731 (2016).

5. Wong, G.J., et al. Frequency, Determinants, and Outcomes of Emboli to Distal and New Territories Related to Mechanical Thrombectomy for Acute Ischemic Stroke. Stroke 52, 2241–2249 (2021).

6. Yemisci, M., et al. Pericyte contraction induced by oxidative-nitrative stress impairs capillary reflow despite successful opening of an occluded cerebral artery. Nat Med 15, 1031–1037 (2009).

7. El Amki, M., et al. Neutrophils Obstructing Brain Capillaries Are a Major Cause of No-Reflow in Ischemic Stroke. Cell Rep 33, 108260 (2020).

8. Stoll, G. & Pham, M. Beyond recanalization - a call for action in acute stroke. Nat Rev Neurol 16, 591–592 (2020).

9. Liebeskind, D.S. Collateral circulation. Stroke 34, 2279–2284 (2003).

10. Schaper, W. Collateral circulation: past and present. Basic Res Cardiol 104, 5–21 (2009).

11. Romero, J.R., et al. Cerebral collateral circulation in carotid artery disease. Curr Cardiol Rev 5, 279–288 (2009).

12. Faber, J.E., Storz, J.F., Cheviron, Z.A. & Zhang, H. High-altitude rodents have abundant collaterals that protect against tissue injury after cerebral, coronary and peripheral artery occlusion. J Cereb Blood Flow Metab 41, 731–744 (2021).

13. Liebeskind, D.S., et al. Collateral Circulation in Thrombectomy for Stroke After 6 to 24 Hours in the DAWN Trial. Stroke 53, 742–748 (2022).

14. Garcia-Tornel, A., et al. Leptomeningeal Collateral Flow Modifies Endovascular Treatment Efficacy on Large-Vessel Occlusion Strokes. Stroke 52, 299–303 (2021).

15. Schuler, F., et al. Differential Benefit of Collaterals for Stroke Patients Treated with Thrombolysis or Supportive Care : A Propensity Score Matched Analysis. Clin Neuroradiol 30, 525–533 (2020).

16. Lucitti, J.L., et al. Variants of Rab GTPase-Effector Binding Protein-2 Cause Variation in the Collateral Circulation and Severity of Stroke. Stroke 47, 3022–3031 (2016).

17. Zhang, H., Prabhakar, P., Sealock, R. & Faber, J.E. Wide genetic variation in the native pial collateral circulation is a major determinant of variation in severity of stroke. J Cereb Blood Flow Metab 30, 923–934 (2010).

18. Renier, N., et al. iDISCO: a simple, rapid method to immunolabel large tissue samples for volume imaging. Cell 159, 896–910 (2014).

19. Kirst, C., et al. Mapping the Fine-Scale Organization and Plasticity of the Brain Vasculature. Cell 180, 780–795 e725 (2020).

20. El Amki, M., et al. Experimental modeling of recombinant tissue plasminogen activator effects after ischemic stroke. Exp Neurol 238, 138–144 (2012).

21. Orset, C., et al. Mouse model of in situ thromboembolic stroke and reperfusion. Stroke 38, 2771–2778 (2007).

22. Dunn, A.K., Bolay, H., Moskowitz, M.A. & Boas, D.A. Dynamic imaging of cerebral blood flow using laser speckle. J Cereb Blood Flow Metab 21, 195–201 (2001).

23. Deffieux, T., Demene, C. & Tanter, M. Functional Ultrasound Imaging: A New Imaging Modality for Neuroscience. Neuroscience 474, 110–121 (2021).

24. Zaidat, O.O., et al. Recommendations on angiographic revascularization grading standards for acute ischemic stroke: a consensus statement. Stroke 44, 2650–2663 (2013).

25. Deng, G., et al. Predictors of futile recanalization after endovascular treatment in acute ischemic stroke: a meta-analysis. J Neurointerv Surg (2021).

26. Hossmann, K.A. Viability thresholds and the penumbra of focal ischemia. Ann Neurol 36, 557–565 (1994).

27. Eltzschig, H.K. & Eckle, T. Ischemia and reperfusion--from mechanism to translation. Nat Med 17, 1391–1401 (2011).

28. Schaffer, C.B., et al. Two-photon imaging of cortical surface microvessels reveals a robust redistribution in blood flow after vascular occlusion. PLoS Biol 4, e22 (2006).

29. Ma, J., Ma, Y., Shuaib, A. & Winship, I.R. Improved collateral flow and reduced damage after remote ischemic perconditioning during distal middle cerebral artery occlusion in aged rats. Scientific reports 10, 12392 (2020).

30. Errico, C., et al. Ultrafast ultrasound localization microscopy for deep super-resolution vascular imaging. Nature 527, 499–502 (2015).

31. Renaudin, N., et al. Functional ultrasound localization microscopy reveals brain-wide neurovascular activity on a microscopic scale. Nat Methods 19, 1004–1012 (2022).

32. Heusch, G. & Gersh, B.J. The pathophysiology of acute myocardial infarction and strategies of protection beyond reperfusion: a continual challenge. Eur Heart J 38, 774–784 (2017).

33. Zhou, M., et al. Myocardial Ischemia-Reperfusion Injury: Therapeutics from a Mitochondria-Centric Perspective. Cardiology 146, 781–792 (2021).

34. Yellon, D.M. & Hausenloy, D.J. Myocardial reperfusion injury. N Engl J Med 357, 1121–1135 (2007).

35. Garcia-Prieto, J., et al. Neutrophil stunning by metoprolol reduces infarct size. Nat Commun 8, 14780 (2017).

36. Kloner, R.A., King, K.S. & Harrington, M.G. No-reflow phenomenon in the heart and brain. American journal of physiology. Heart and circulatory physiology 315, H550–H562 (2018).

37. Gauberti, M., et al. Ischemia-Reperfusion Injury After Endovascular Thrombectomy for Ischemic Stroke. Stroke 49, 3071–3074 (2018).

38. Carmichael, S.T. Rodent models of focal stroke: size, mechanism, and purpose. NeuroRx 2, 396–409 (2005).

39. Doyle, K.P. & Buckwalter, M.S. A mouse model of permanent focal ischemia: distal middle cerebral artery occlusion. Methods Mol Biol 1135, 103–110 (2014).

40. Chauveau, F., et al. In vivo MRI assessment of permanent middle cerebral artery occlusion by electrocoagulation: pitfalls of procedure. Exp Transl Stroke Med 2, 4 (2010).

41. Qazi, E.M., et al. Thrombus Characteristics Are Related to Collaterals and Angioarchitecture in Acute Stroke. Can J Neurol Sci 42, 381–388 (2015).

42. Seners, P., et al. Better Collaterals Are Independently Associated With Post-Thrombolysis Recanalization Before Thrombectomy. Stroke 50, 867–872 (2019).

43. Shimonaga, K., et al. Early venous filling after reperfusion therapy in acute ischemic stroke. J Stroke Cerebrovasc Dis 29, 104926 (2020).

44. Zhao, G., et al. Evaluation of the role of susceptibility-weighted imaging in thrombolytic therapy for acute ischemic stroke. Journal of clinical neuroscience : official journal of the Neurosurgical Society of Australasia 40, 175–179 (2017).

45. Bornstein, N.M., et al. An injectable implant to stimulate the sphenopalatine ganglion for treatment of acute ischaemic stroke up to 24 h from onset (ImpACT-24B): an international, randomised, double-blind, sham-controlled, pivotal trial. Lancet 394, 219–229 (2019).

46. Holtmaat, A., et al. Long-term, high-resolution imaging in the mouse neocortex through a chronic cranial window. Nat Protoc 4, 1128–1144 (2009).

47. El Amki, M., et al. Improved Reperfusion and Vasculoprotection by the Poly(ADP-Ribose)Polymerase Inhibitor PJ34 After Stroke and Thrombolysis in Mice. Molecular neurobiology 55, 9156–9168 (2018).

48. Sekhon, L.H., Spence, I., Morgan, M.K. & Weber, N.C. Chronic cerebral hypoperfusion in the rat: temporal delineation of effects and the in vitro ischemic threshold. Brain Res 704, 107–111 (1995).

49. Maysami, S., et al. A cross-laboratory preclinical study on the effectiveness of interleukin-1 receptor antagonist in stroke. J Cereb Blood Flow Metab 36, 596–605 (2016).

50. Mayrhofer, J.M., et al. Design and performance of an ultra-flexible two-photon microscope for in vivo research. Biomed Opt Express 6, 4228–4237 (2015).

51. Pologruto, T.A., Sabatini, B.L. & Svoboda, K. ScanImage: flexible software for operating laser scanning microscopes. Biomed Eng Online 2, 13 (2003).

52. Nouhoum, M., et al. A functional ultrasound brain GPS for automatic vascular-based neuronavigation. Scientific reports 11, 15197 (2021).

53. Voigt, F.F., et al. The mesoSPIM initiative: open-source light-sheet microscopes for imaging cleared tissue. Nat Methods 16, 1105–1108 (2019).

54. El Amki, M., et al. Contraceptive drugs mitigate experimental stroke-induced brain injury. Cardiovascular research 115, 637–646 (2019).

55. Haddad, M., et al. Reduction of hemorrhagic transformation by PJ34, a poly(ADP-ribose)polymerase inhibitor, after permanent focal cerebral ischemia in mice. Eur J Pharmacol 588, 52–57 (2008).

56. Wahl, F., Allix, M., Plotkine, M. & Boulu, R.G. Neurological and behavioral outcomes of focal cerebral ischemia in rats. Stroke 23, 267–272 (1992).

57. Lyden, P., et al. Improved reliability of the NIH Stroke Scale using video training. NINDS TPA Stroke Study Group. Stroke 25, 2220–2226 (1994).

58. Banks, J.L. & Marotta, C.A. Outcomes validity and reliability of the modified Rankin scale: implications for stroke clinical trials: a literature review and synthesis. Stroke 38, 1091–1096 (2007).

59. Campbell, B.C., et al. Endovascular therapy for ischemic stroke with perfusion-imaging selection. N Engl J Med 372, 1009–1018 (2015).

